# Optineurin provides a mitophagy contact site for TBK1 activation

**DOI:** 10.1101/2023.02.24.529790

**Authors:** Koji Yamano, Momoha Sawada, Reika Kikuchi, Kafu Nagataki, Waka Kojima, Atsushi Sugihara, Tomoshige Fujino, Keiji Tanaka, Gosuke Hayashi, Hiroshi Murakami, Noriyuki Matsuda

## Abstract

Tank-binding kinase 1 (TBK1) is a Ser/Thr kinase involved in many intracellular processes including innate immunity, cell cycle, and apoptosis. TBK1 is also important for phosphorylating autophagy adaptors critical in selective autophagic removal of damaged mitochondria (mitophagy). However, the mechanism by which TBK1 is activated by PINK1/Parkin-mediated mitophagy remains largely unknown. Here, we show that the autophagy adaptor OPTN provides a unique platform for TBK1 activation. The OPTN-ubiquitin and OPTN-autophagy machinery interaction axes facilitate assembly of the OPTN-TBK1 complex at a contact site between damaged mitochondria and the autophagosome formation site. This assembly point serves as a positive feedback loop for TBK1 activation by accelerating hetero-autophosphorylation of the protein. Furthermore, expression of monobodies engineered in this study against OPTN impaired assembly of OPTN at the contact sites as well as the subsequent activation of TBK1 and mitochondrial degradation. Taken together the findings reveal that a positive reciprocal relationship between OPTN and TBK1 initiates autophagosome biogenesis on damaged mitochondria.

## Introduction

Macroautophagy (hereafter referred to as autophagy) is a conserved catabolic process that delivers cytoplasmic components, organelles and cellular pathogens to lysosomes for degradation. These cargo/substrates are encapsulated by double membrane structures called autophagosomes that subsequently fuse with lysosomes for cargo degradation. Various proteins have been identified as drivers of the *de novo* autophagosomal membrane synthesis. Although autophagosome biogenesis is induced by various signals including starvation, oxidative stress and DNA damage, the unique process of autophagosome formation begins with ATG9A-containing vesicles and the ULK complex, which in mammals is comprised of FIP200, ATG13, ATG101 and ULK1/2 kinases. ATG9A functions as a lipid scramblase and the ATG9A-containing vesicles act as initial isolation membrane seeds for autophagosome formation ^1–3^. Incorporation of multiple ULK1 complexes into the developing scaffold further contributes to the recruitment of downstream autophagy proteins ^4, 5^. Autophosphorylation of ULK1/2 in turn triggers ULK1/2-dependent phosphorylation of diverse downstream substrates including ATG9A, BECN1 and ATG14 in mammals ^6–10^. Subsequent activation of a phosphatidylinositol 3-kinase complex composed of BECN1, ATG14, VPS15 and VPS34 is required for the production of the phosphatidylinositol 3-phosphate that facilitates recruitment of the ATG2-WIPI complex to the autophagosomal formation site ^11^. Two ubiquitin-like conjugation systems, the ATG5-ATG12/ATG16L1 complex and phosphatidylethanolamine-conjugated ATG8 (LC3 family proteins in mammals), are important for elongation of the isolation membrane as well as efficient degradation of the inner autophagosomal membrane in lysosomes ^12–14^.

In addition to the autophagy core kinase units (the ULK and PI3K complexes), other phosphorylation systems regulate autophagy initiation and/or progression. Under nutrient-rich conditions, mammalian TORC1 (mTORC1) suppresses activity of the ULK complex via ULK1/2 and ATG13 phosphorylation ^15, 16^. Furthermore, during Parkin-driven mitochondrial autophagy (mitophagy), the kinases PINK1 and TBK1 are important for Parkin activation and autophagosome formation, respectively (see further details later).

Although autophagy has been recognized as a bulk degradation process that non-selectively degrades cytoplasmic components, multiple lines of evidence show that selective-type autophagy mediates the clearance of specific unwanted/damaged organelles ^17^. PINK1/Parkin-mediated mitophagy is one of the best characterized selective pathways that functions to maintain cellular homeostasis ^18–20^. Defects in mitophagy have been linked to neurodegeneration in Parkinson’s disease. The E3 ligase Parkin and the mitochondrial kinase PINK1, both of which are causal gene products of familial Parkinson disease ^21, 22^, coordinately function to generate poly-ubiquitin (Ub) chains on damaged mitochondria. Under healthy mitochondrial conditions, PINK1 is constitutively and rapidly degraded by proteasomes after cleaving the mitochondrial targeting sequence and the associated hydrophobic transmembrane segments in the mitochondria ^23–29^. In contrast, when mitochondria are damaged by aging and/or exposure to toxic compounds and lose their membrane potential, PINK1 accumulates on the outer membrane via associations with the TOMM complex ^30–32^. PINK1 kinase activity is essential for Parkin recruitment to damaged mitochondria as well as activation of Parkin E3 activity ^33–35^. PINK1 phosphorylates Ub and the Ub-like (UBL) domain of Parkin at Serine 65 ^36–40^. Binding phosphorylated Ub and UBL phosphorylation promote conformational changes in the Parkin structure that convert the inactivated form to the activated form ^41–44^. Consequently, PINK1, Parkin, Ub, and various E2 enzymes form a positive feedback loop of Ub amplification on damaged mitochondria ^45–48^. Ub-coated mitochondria are then recognized by autophagy adaptors ^49, 50^. Autophagy adaptors (also known as autophagy receptors) are defined as proteins containing both a Ub-binding domain and an LC3 interacting region (LIR), with five different adaptors, OPTN, NDP52, TAX1BP1, p62 and NBR1, identified to date in mammals ^51^. Autophagy adaptors thus function as a molecular bridge linking ubiquitinated cargo with the autophagy machinery. In PINK1/Parkin-mediated mitophagy, OPTN and NDP52 are required for autophagosome formation ^49^. NDP52 is composed of several functional domains such as SKICH for interactions with AZI2 and TBK1BP1, LIR for LC3 binding, coiled-coil for homo-dimerization, and two C-terminal zinc fingers for interactions with the Ub-chain ^50^. In addition to LC3 binding, NDP52 also directly binds FIP200, which promotes recruitment of the ULK complex to damaged mitochondria ^52–55^.

OPTN is another essential mitophagy adaptor composed of long coiled-coil regions, a leucin zipper and an LIR as well as UBAN and zinc finger domains ^50^. Recent experimental evidence has shown that OPTN interacts with multiple autophagy components during Parkin-mediated mitophagy. In addition to LC3 binding via the LIR, OPTN can recruit ATG9A vesicles through its leucine zipper motif ^56^. Furthermore, the LIR in OPTN can interact with FIP200 when OPTN S177 is phosphorylated ^57^. Given the multiple interactions linking NDP52 and OPTN with autophagy components, they are essential adaptors for PINK1/Parkin-mediated mitophagy.

In addition to autophagy adaptors, TBK1-mediated phosphorylation regulates Ub-dependent selective autophagy ^58^. TBK1 interacts directly with OPTN and indirectly with NDP52 and TAX1BP1 via AZI2 and TBKBP1 ^59, 60^. During mitophagy, autophosphorylation of TBK1 at S172 converts the protein to its activated form ^61, 62^. Activated TBK1 subsequently phosphorylates a number of autophagy adaptors ^63^. Furthermore, it also phosphorylates RAB7A, which promotes ATG9A recruitment to damaged mitochondria ^64^, and LC3C and GABARAPL2 that expand the isolation membranes ^65^. Interestingly, gene mutations in both *OPTN* and *TBK1* have been found in patients suffering from amyotrophic lateral sclerosis (ALS) and frontotemporal dementia (FTD), suggesting physiological and molecular links between OPTN and TBK1 ^66, 67^. While TBK1 autophosphorylation is a prerequisite for TBK1 activation, the mechanism underlying this mitophagy-driven step remains unknown.

In this study, we found that *TBK1* deletion inhibits the localization of OPTN, but not NDP52, at the autophagosome formation site during mitophagy, and correspondingly that deletion of *OPTN*, but not *NDP52*, inhibits TBK1 autophosphorylation. We also found that activated TBK1 is selectively and rapidly degraded via autophagy. Two different interaction axes, OPTN-Ub and OPTN-autophagy, enable the formation of a contact site between the isolation membrane and damaged mitochondria that supports autophosphorylation of TBK1 by neighboring TBK1. In addition, monobodies engineered in this study against OPTN inhibited formation of the OPTN-dependent contact site, which suppressed both TBK1 autophosphorylation and the elimination of damaged mitochondria. These results indicate that OPTN-TBK1 catalyzes a unique autophagy initiation step that is different from that mediated by NDP52.

## Results

### OPTN is required for TBK1 phosphorylation and subsequent autophagic degradation.

Although autophosphorylation of TBK1 at S172 was reported to increase during PINK1/Parkin-mediated mitophagy ^62^, how it is regulated remains largely unknown. Immunoblots of HeLa cells stably expressing Parkin and which had been treated with valinomycin for varying lengths of time revealed phosphorylation of S172 in TBK1. As shown in Fig 1a, this phosphorylation event was observed after 30 min valinomycin treatment, with the levels gradually decreasing over time. While phosphatase-mediated de-phosphorylation has been suggested for the OPTN-TBK1 axis ^68^, we found that reduction of the TBK1 pS172 signal was blocked when cells were treated with the lysosomal inhibitor bafilomycin A1 (Fig 1a, b). Concanamycin A, a different lysosomal inhibitor, similarly enhanced the TBK1 pS172 signal, whereas the proteasomal inhibitors epoxomicin and MG132 had no effect (Fig 1c, d), indicating that the TBK1 phosphorylation induced by Parkin-mediated ubiquitination is rapidly degraded by autophagy pathway in lysosomes. Consequently, to limit autophagic degradation of activated TBK1 during PINK1/Parkin-mediated mitophagy, we used bafilomycin A1 for subsequent analyses. TBK1 phosphorylates S177 in OPTN, which undergoes ubiquitination and autophagic degradation (Fig 1a, b). Since OPTN and NDP52 are essential adaptors for mitophagy and TBK1 is either directly (OPTN) or indirectly (NDP52) associated with both, we next examined the requirement of the two adaptors for TBK1 autophosphorylation. Although *OPTN* KO drastically reduced the level of phosphorylated TBK1 at S172, *NDP52* KO effects on the pS172 signal were comparable to controls (Fig 1e, f). Similar results were observed when OPTN and NDP52 were re-expressed in Penta KO HeLa cells lacking all five autophagy adaptors (Fig 1g). Furthermore, there was a significant increase in the TBK1 pS172 signal in cells overexpressing OPTN and which had been treated with valinomycin and bafilomycin A1 for 3 hrs as compared to cells that lacked OPTN overexpression (Fig 1h, i). These results indicate that OPTN is a crucial rate-limiting factor for the autophosphorylation of TBK1 that occurs in response to PINK1/Parkin-mediated mitophagy.

**Figure 1.**
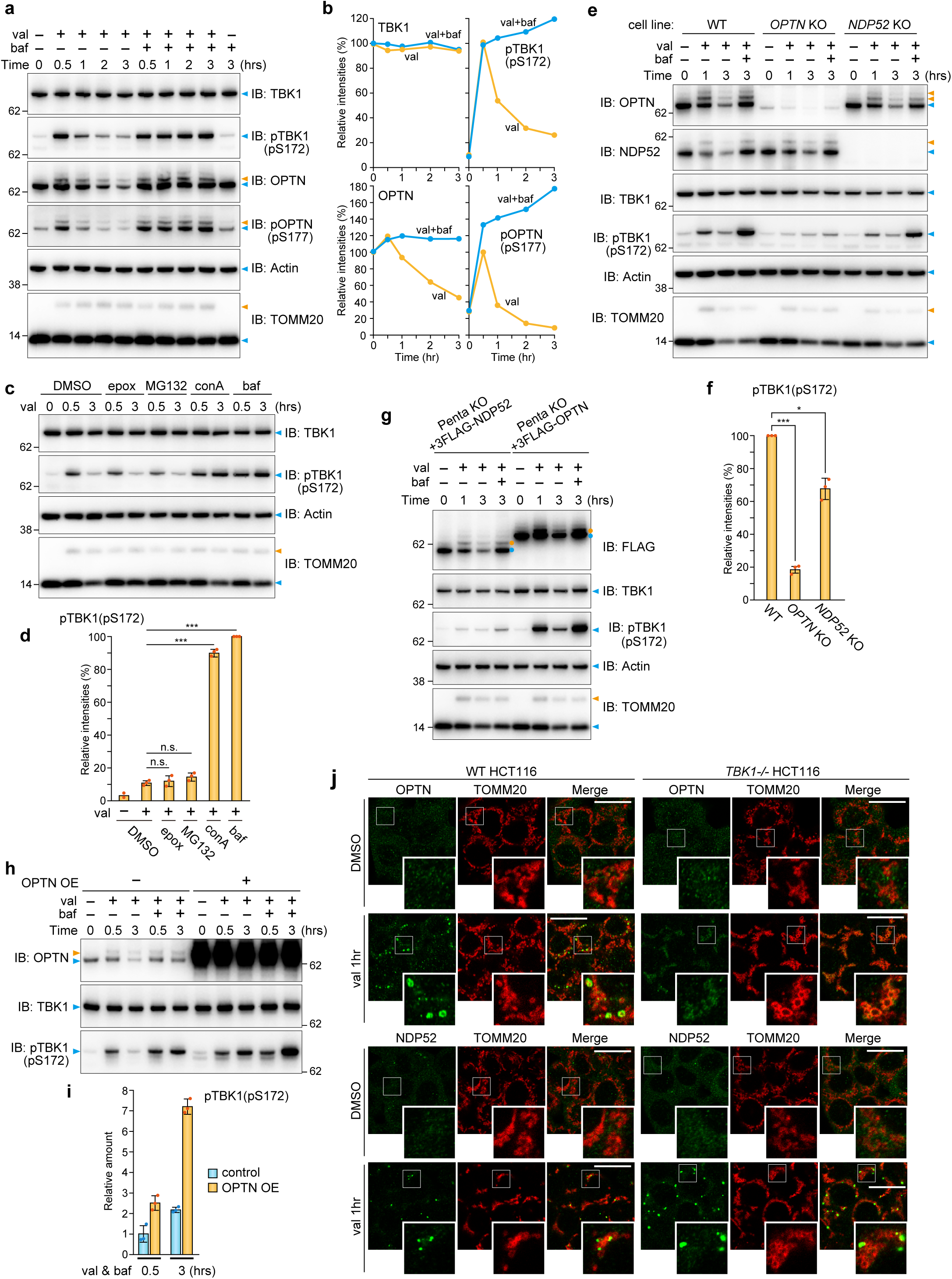
Activated TBK1 induced by OPTN during mitophagy is rapidly degraded via autophagy. **a** HeLa cells stably expressing Parkin were treated with valinomycin (val) and bafilomycin (baf) for the indicated times. Total cell lysates were analyzed by immunoblotting (IB). Orange arrowheads indicate ubiquitinated bands. **b** The levels of proteins indicated in (a) were quantified. TBK and OPTN levels without valinomycin were set to 100. pTBK1(pS172) and pOPTN(pS177) levels after 0.5 hrs val were set to 100. **c** Parkin-expressing HeLa cells treated with val and the indicated inhibitors were analyzed by IB. Epox; Epoxomicin, conA; Concanamycin A. **d** pTBK1 levels after 3 hrs val treatment in (c) were quantified. Error bars represent mean ± s.d. of three independent experiments. **e** WT, *OPTN* KO, and *NDP52* KO HeLa cells stably expressing Parkin were treated with val and baf for the indicated times. Total cell lysates were analyzed by IB. Orange arrowheads indicate ubiquitinated bands. **f** pTBK1 levels after 3 hrs val and baf treatment in (e) were quantified. Error bars represent mean ± s.d. of three independent experiments. **g** Penta KO HeLa cells stably expressing Parkin and 3FLAG-NDP52 or 3FLAG-OPTN were treated with val and baf for the indicated times and analyzed by IB. Orange arrowheads indicate ubiquitinated bands. **h** Parkin-expressing HeLa cells with or without 3FLAG-OPTN overexpression (OE) were treated with val and baf for the indicated times and analyzed by IB. Orange arrowheads indicate ubiquitinated bands. **i** pTBK1 levels in Parkin-expressing HeLa cells with or without OPTN OE in (h) were quantified. Error bars represent mean ± s.d. of three independent experiments. pTBK1 levels in control cells after 0.5 hrs val and baf treatment were set to 1. **j** WT and *TBK1-/-* HCT116 cells stably expressing Parkin were treated with DMSO or val for 1 hr and then immunostained with the indicated antibodies. Bars, 10 μm.

We next examined the recruitment of OPTN and NDP52 to damaged mitochondria in cells lacking TBK1. In WT cells, both OPTN and NDP52 formed cup-shaped and/or sphere-like structures on damaged mitochondria (Fig 1j) that were previously reported to co-localize with LC3B-labeled autophagic membranes ^49, 56, 69^. In *TBK1-/-* HCT116 cells, the NDP52 structures were larger with more pronounced signals (Fig 1j). In sharp contrast, OPTN was dispersed throughout the damaged mitochondria rather than being localized in concentrated spots (Fig 1j). Thus, although NDP52 and OPTN are essential mitophagy-adaptors that interact with TBK1, their recruitment to the autophagosome formation site is differently regulated by TBK1.

We also observed mitophagy-dependent recruitment of TBK1, as well as other autophagy adaptors, to damaged mitochondria at endogenous expression levels. HeLa cells stably expressing Parkin were treated with valinomycin for 1 hr and then immunostained. While the endogenous TBK1 signal could not be observed using commercially available antibodies, the TBK1 pS172 signal was detected on damaged mitochondria (Supplementary Fig 1a). Spherical/cup-shaped and dot-like signals were observed at the periphery of the mitochondria (Supplementary Fig 1a). Similar signals were observed using an anti-pS177 OPTN antibody (Supplementary Fig 1a). In contrast, the TAX1BP1 signal was dispersed throughout the damaged mitochondria (Supplementary Fig 1a). Although p62 and NBR1 formed dot-like structures on the mitochondria, they had clearly accumulated at contact sites linking two mitochondria (Supplementary Fig 1a). Consistent with these findings, phosphorylated TBK1, OPTN, and NDP52 also colocalized with GFP-LC3B in response to PINK1/Parkin-mediated mitophagy, whereas p62 did not (Supplementary Fig 1b).

### OPTN association with the autophagy machinery is required for TBK1 activation

We next investigated how OPTN induces TBK1 autophosphorylation during mitophagy. Previously, OPTN binding of Ub was shown to be a requirement for TBK1 autophosphorylation ^62^. Here, we found that OPTN association with the autophagy machinery is also required for TBK1 activation. The OPTN leucine zipper and LIR motifs are important for the recruitment of ATG9A vesicles, FIP200, and LC3 family proteins as 4LA mutations in the leucine zipper motif disrupted ATG9A vesicle interactions ^56^ and an F178A mutation in the LIR inhibited FIP200/LC3 family protein interactions ^57, 63, 70^. Penta KO HeLa cells stably expressing OPTN WT or the previously defined OPTN mutations along with Parkin were treated with valinomycin and the phosphorylation state of TBK1 was assessed. TBK1 phosphorylation, however, was not apparent in the OPTN mutant lines, even after 3 hrs with valinomycin, indicating that autophagy adaptors are essential for TBK1 activation (Fig 2a). OPTN WT expression in Penta KO cells induced autophosphorylation of TBK1 at S172 after 1 hr valinomycin treatment (Fig 2a). This signal was further enhanced with bafilomycin A1 (Fig 2a). In contrast, the OPTN 4LA and F178A mutants as well as the double 4LA/F178A mutant had decreased levels of phosphorylated TBK1 compared to cells expressing OPTN WT (Fig 2a, b). When the intracellular localization was observed, OPTN WT signals were visible as cup-shaped and/or sphere-like structures on the mitochondria (Fig 2c) that colocalized with isolation membranes labelled by LC3B (Supplementary Fig 1b). Despite clear association with damaged mitochondria, recruitment of OPTN to the autophagosome formation site was reduced in the F178A mutant and was abolished by the 4LA and 4LA/F178A double mutants (Fig 2c). These results indicate that association of OPTN with damaged mitochondria alone is not sufficient, and that recruitment of OPTN to the autophagosome formation site, where isolation membranes are synthesized and expanded, is required for full activation of TBK1.

**Figure 2.**
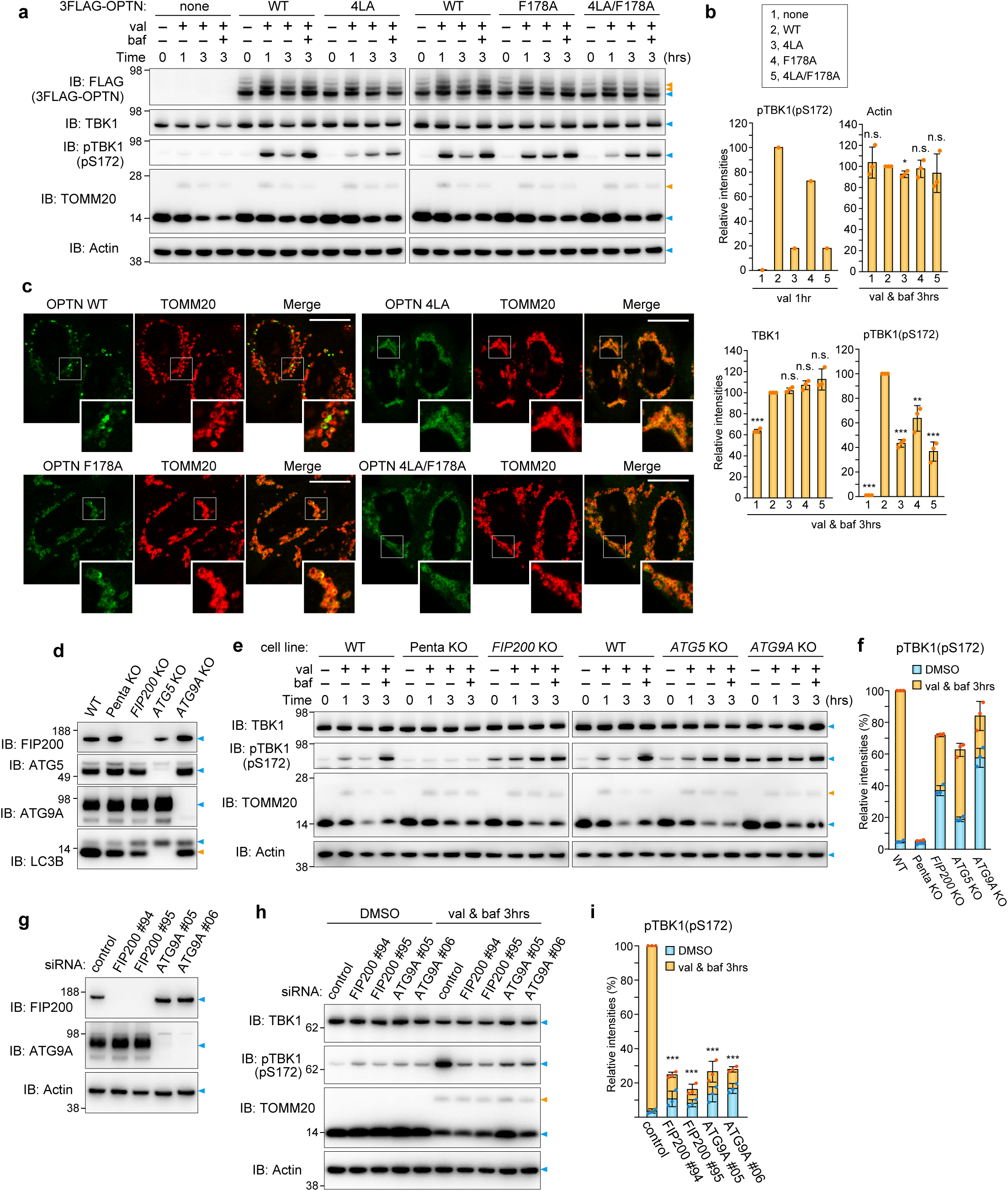
TBK1 activation requires OPTN-autophagic machinery associations. **a** Penta KO HeLa cells stably co-expressing Parkin and 3FLAG-OPTN WT or the indicated mutants were treated with valinomycin (val) and bafilomycin (baf) for the indicated times. Total cell lysates were analyzed by immunoblotting (IB). Orange arrowheads indicate ubiquitinated bands. **b** The levels of proteins indicated in (a) were quantified. The protein levels in cells expressing OPTN WT were set to 100. Error bars represent mean ± s.d. of three independent experiments. **c** Cells treated with val for 3 hrs in (a) were immunostained with the indicated antibodies. Bars, 10 μm. **d** Depletion of the indicated proteins was confirmed by IB. The orange arrowhead denotes lipidated forms of LC3B. **e** The indicated cells stably expressing Parkin were treated with val and baf for the indicated times. Total cell lysates were analyzed by IB. Orange arrowheads indicate ubiquitinated bands. **f** The levels of pTBK1 in (e) were quantified. The pTBK1 level in WT cells treated with val and bad for 3 hrs was set to 100. Error bars represent mean ± s.d. of three independent experiments. **g** HeLa cells stably expressing Parkin treated with the indicated siRNA were analyzed by IB. **h** HeLa cells stably expressing Parkin pre-treated with the indicated siRNA were treated with val and baf for 3 hrs and analyzed by IB. Orange arrowheads indicate ubiquitinated bands. **i** The levels of pTBK1 in (h) were quantified. The pTBK1 level in control cells was set to 100%. Error bars represent mean ± s.d. of three independent experiments.

Next, to confirm that autophagy machinery is required for TBK1 activation during mitophagy, autophagy gene KO (*FIP200* KO, *ATG5* KO and *ATG9A* KO) cells were used (Fig 2d). The levels of FIP200, ATG5, ATG9A, and lipidated LC3B in Penta KO cells are comparable to those in WT cells (Fig 2d). When Parkin-mediated ubiquitination was induced, phosphorylation of TBK1 at S172 was observed in WT cells after 1 hr valinomycin and was significantly enhanced after 3 hrs with bafilomycin (Fig 2e). Although autophagy gene KO cells possess a certain amount of phosphorylated TBK1 even under basal growing conditions (*i.e.,* valinomycin 0 hr), the levels of TBK1 pS172 did not increase during mitophagy (Fig 2e). To measure the amounts of newly generated TBK1 pS172, we subtracted the basal phosphorylation signal from that generated post-valinomycin (1 hr) and bafilomycin (3 hr). In WT cells, the phosphorylation signal was ∼ 90 but was less than 30 in *ATG9A* KO cells (Fig 2f). Thus, although phosphorylated TBK1 accumulates in autophagy gene KO cells, TBK1 pS172 generated in response to Parkin-mediated ubiquitination requires the autophagy machinery. Further, FIP200 and ATG9A, rather than ATG5, are critical for mitophagy-dependent phosphorylation of TBK1, suggesting that the initial step in isolation membrane biogenesis is more important for phosphorylation. Autophagy gene KOs constitutively inhibit autophagy, thereby causing an accumulation of phosphorylated TBK1 under normal growing conditions (Fig 2e). To simultaneously inhibit autophagy and induce Parkin-mediated ubiquitination, we knocked down FIP200 and ATG9A by siRNA before inducing mitochondrial damage. As shown in Fig 2g, both siRNAs efficiently knocked down FIP200 and ATG9A. While there was a slight increase in TBK1 phosphorylation in the FIP200 and ATG9A siRNA-treated cells under basal conditions, the signal following Parkin-mediated ubiquitination was only moderately elevated when compared to that in control cells (Fig 2h, i). These results demonstrate that both ubiquitination and the autophagy core subunits are required for TBK1 autophosphorylation of S172 during mitophagy.

### TBK1 activation does not require OPTN under basal autophagy conditions

Because cells with autophagy gene deletions accumulate phosphorylated TBK1 (Fig 2e), we next examined whether OPTN mediates TBK1 autophosphorylation under basal conditions. Protein levels for the autophagy adaptors NDP52, TAX1BP1, p62, and NBR1 all increased following KO of the autophagy genes *FIP200*, *ATG5* and *ATG9A* (Supplementary Fig 2), whereas OPTN levels remained constant (Supplementary Fig 2). Further, TBK1BP1, the mediator between TBK1 and NDP52/TAX1BP1, became unstable in Penta KO cells and the electrophoretic migration of AZI2, another TBK1 mediator, was altered following deletion of either *FIP200* or *ATG9A* (Supplementary Fig 2). Although total TBK1 levels were comparable between WT and autophagy gene KO cells, TBK1 pS172 levels were moderately increased in *ATG5* KO cells and significantly elevated in *FIP200* KO and *ATG9A* KO cells (Supplementary Fig 2). p62 (S403) phosphorylation was also elevated (Supplementary Fig 2). We next examined the subcellular localization pattern of autophagy adaptors in the KO cells. NDP52, TAX1BP1, p62, and NBR1 as well as phosphorylated TBK1 colocalized with Ub-positive condensates in *FIP200* KO and *ATG9A* KO cells (Fig 3a). Ferritin was also concentrated on the p62-positive condensates (Fig 3a). In contrast, most of the OPTN signal was cytosolic and did not colocalize with the Ub condensates (Fig 3a).

**Figure 3.**
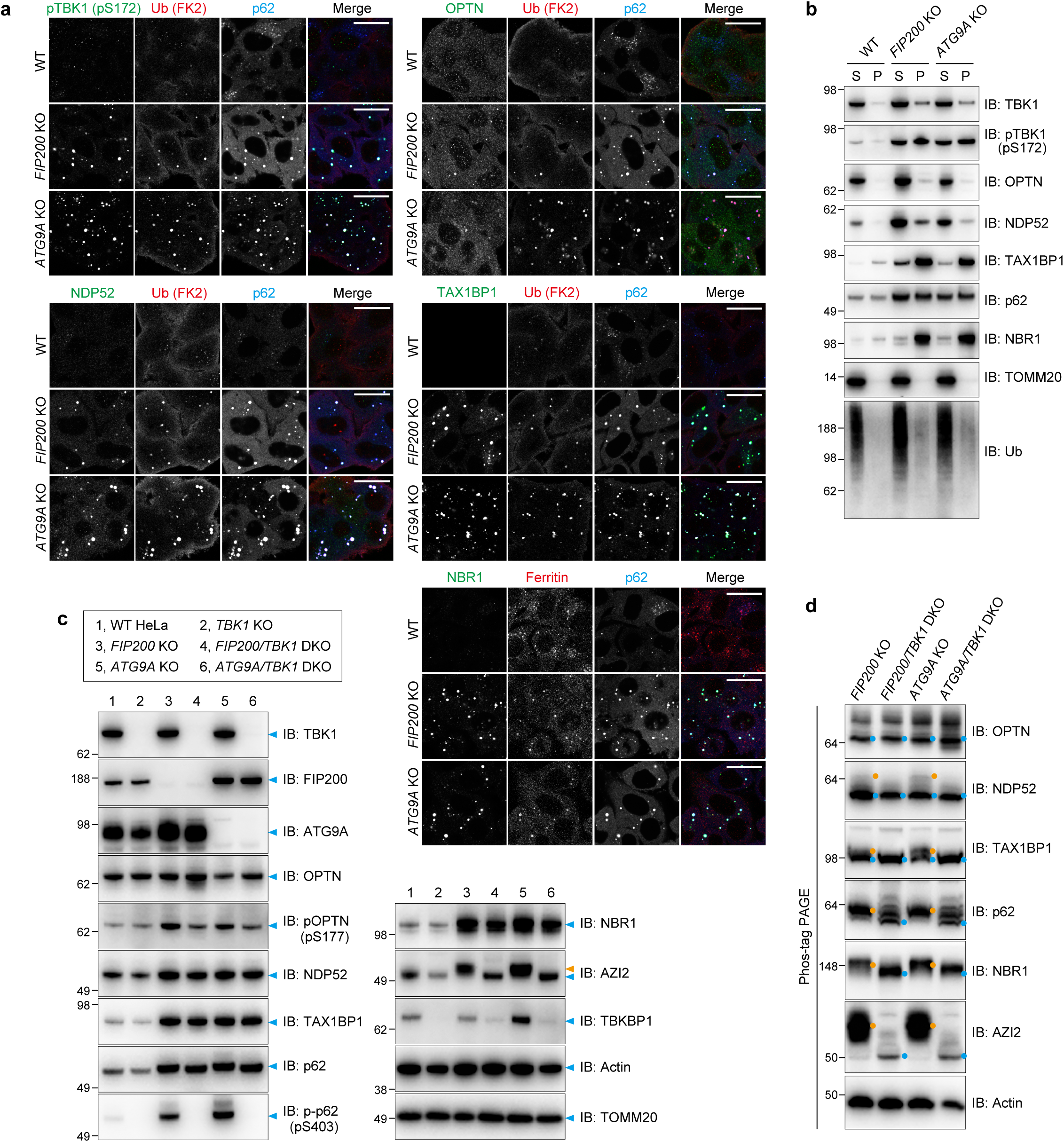
TBK1 that accumulates in autophagy gene KO cells phosphorylates autophagy adaptors other than OPTN. **a** WT, *FIP200* KO, and *ATG9A* KO HeLa cells were immunostained with the indicated antibodies. **b** WT, *FIP200* KO, and *ATG9A* KO HeLa cells were solubilized using 2% TritonX-100 with the supernatants (S) and pellets (P) separated by centrifugation and then analyzed by immunoblotting (IB). **c** The indicated proteins in WT, *TBK1* KO, *FIP200* KO, *FIP200/TBK1* DKO, *ATG9A* KO, and *ATG9A/TBK1* DKO HeLa cells were analyzed by IB. The orange arrowhead indicates a slower migrating AZI2. **d** Total cell lysates prepared in (c) were analyzed by Phos-tag PAGE followed by IB. Orange dots indicate proteins phosphorylated by TBK1.

Biochemical analyses also showed that insoluble forms of the autophagy adaptors were elevated following autophagy gene depletion (Fig 3b). After solubilizing WT, *FIP200* KO, and *ATG9* KO cells with 2% TX-100, the soluble and insoluble pellet fractions were separated by centrifugation. We found that autophagy KO induced insoluble Ub smears and an accumulation of NDP52, TAX1BP1, p62, and NBR1 in the insoluble fractions (Fig 3b). Although total protein levels of TBK1 did not change with autophagy KO, a small amount of TBK1 was found in the insoluble fraction (Fig 3b). In addition, the recovery ratio of phosphorylated TBK1 in the insoluble fraction was significantly higher than that of total TBK1 (Fig 3b), indicating that the phosphorylated form is specifically targeted to the insoluble fraction. However, a large percentage of OPTN was maintained in the soluble fraction even in the autophagy KO cells (Fig 3b). Therefore, these results strongly suggest that autophagy adaptors other than OPTN are recruited to Ub-condensates and mediate TBK1 autophosphorylation during basal autophagy conditions. The absence of OPTN recruitment to Ub condensates may be attributable to the type of ubiquitin chain linkages that accumulate following autophagy gene KO ^71^. OPTN-Ub interactions depend on the UBAN domain, which favors linear and K63-linked Ub chains rather than the K48-linked chains that dominate when autophagy pathways are impaired.

Since phosphorylated TBK1 accumulates with autophagy adaptors in Ub-condensates, it may induce phosphorylation of the adaptor during basal autophagy pathway. To test this possibility, we knocked out *TBK1* in WT, *FIP200* KO, and *ATG9A* KO cells (Fig 3c). Although autophagy gene depletion promoted an increase in NDP52, TAX1BP1, p62, and NBR1, their levels were not affected with the concomitant deletion of *TBK1*. However, it completely impeded p62 (S403) phosphorylation and altered the electrophoretic migration AZI2. To clarify the phosphorylation state of other autophagy adaptors/mediators, we next performed Phos-tag analysis (Fig 3d). Although electrophoretic migration of OPTN was not affected by *TBK1* deletion, the other autophagy adaptors in the *FIP200* and *ATG9A* KO cells resolved as several distinct bands and slower migrating bands (denoted by orange-colored dots) disappeared and/or were only present in their unphosphorylated state (denoted by blue-colored dots) (Fig 3d). These results strongly suggest that NDP52, TAX1BP1, p62, NBR1, and AZI2 form Ub-positive condensates where TBK1 is activated, and that activated TBK1 phosphorylates these proteins under basal autophagy conditions, whereas OPTN-mediated activation of TBK1 is specific to Parkin-mediated mitophagy.

### Engineered OPTN-Ub condensates induce TBK1 activation

We next sought to engineer OPTN-Ub condensates and determine if they induce TBK1 activation. For this purpose, we used the fluoppi technique (Fig 4a). Fluoppi can create LLPS (liquid-liquid phase separation) via multimeric interactions ^42, 72^. An HA-Ash tag that forms a homo-oligomer was fused to a linear six-tandem Ub chain to yield HA-Ash-6Ub, and a homotetrametic humanized Azami-Green (hAG) was fused to OPTN to make hAG-OPTN. Co-expression of HA-Ash-6Ub and hAG-OPTN in cells induces multimeric interactions (Ash-Ash, hAG-hAG, and OPTN-Ub) that result in the formation of fluoppi foci in the cytosol (Fig 4a). We and others previously showed that TBK1 rather than the ULK complex is recruited to OPTN-Ub fluoppi foci ^56^ and that M44Q/L5Q mutations in OPTN impede OPTN-TBK1 interactions ^59^. Here, we found that phosphorylation of OPTN (S177) and TBK1 (S172) was induced by the OPTN-Ub fluoppi (Fig 4b) and that neither OPTN nor TBK1 were phosphorylated following *TBK1* deletion (*TBK1-/-*) (Fig 4b). Further, signals for phosphorylated TBK1 (S172) and OPTN (S177) were exclusively localized in the foci (Fig 4c, d), suggesting that TBK1 autophosphorylation occurs in the OPTN condensates. Although OPTN-Ub foci formation still occurred in *TBK1-/-* cells, the phosphorylated OPTN (S177) signal was absent (Fig 4c-f). These results indicate that OPTN can induce TBK1 autophosphorylation when multimeric OPTNs are sequestered within a particular cellular localization. In other words, Parkin-mediated mitophagy promotes interactions between OPTN and the autophagy components that contribute to OPTN multimerization and concomitant TBK1 autophosphorylation.

**Figure 4.**
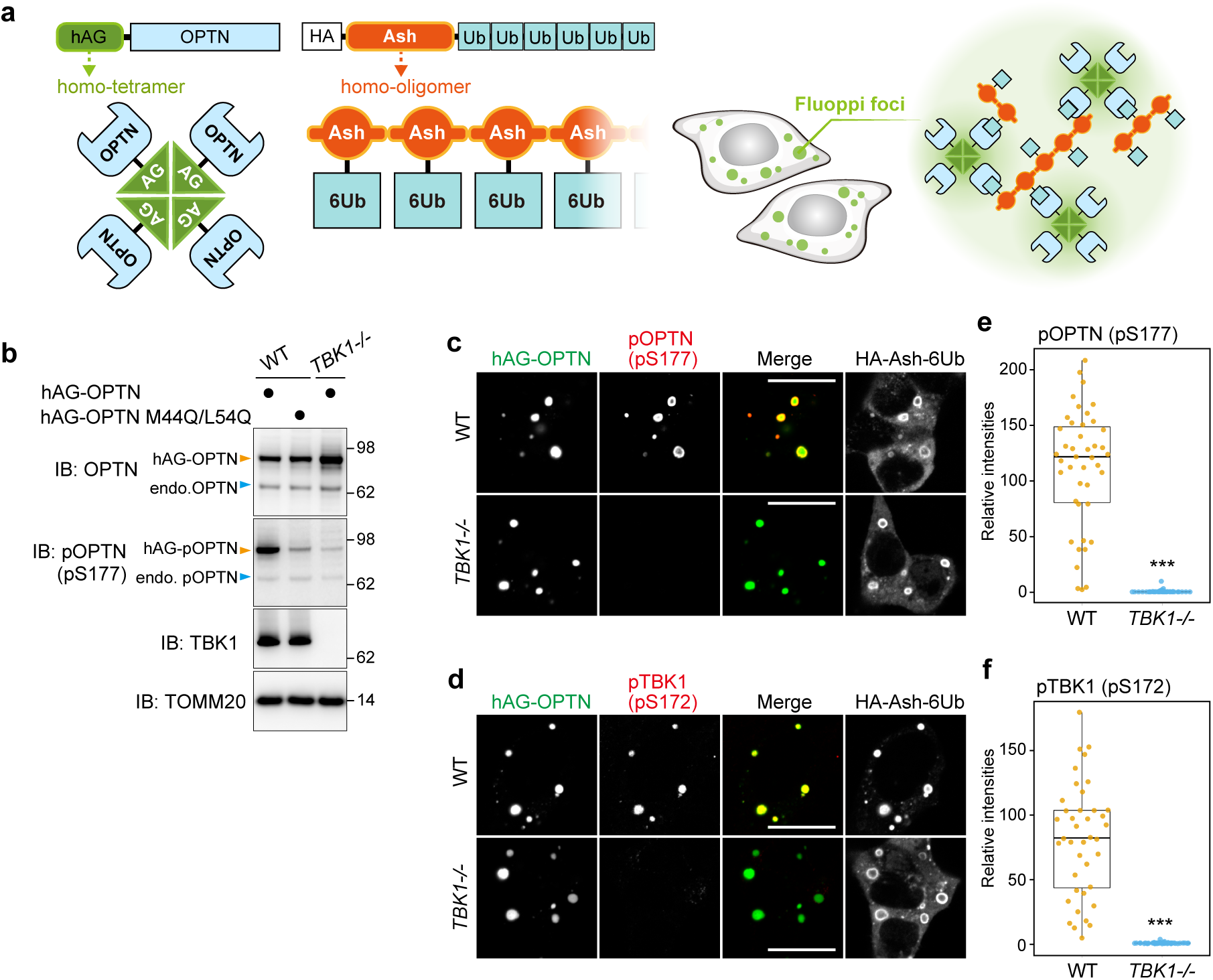
OPTN-Ub condensates created by fluoppi activate TBK1. **a** Schematic representation of fluoppi foci. **b** The indicated hAG-OPTN constructs were expressed with HA-Ash-6Ub in WT and *TBK1-/-* HCT116 cells. Total cell lysates were analyzed by immunoblotting. **c** and **d** hAG-OPTN and HA-Ash-6Ub were expressed in WT and *TBK1-/-* HCT116 cells. The cells were immunostained with antibodies against HA, pOPTN(pS177) for (c) and pTBK1(pS172) for (d). Bars, 10 μm. **e** and **f** The intensities of pOPTN(pS177) and pTBK1(pS172) in the fluoppi foci in (c) and (d), respectively, were quantified. Error bars represent mean ± s.d. with 50 fluoppi foci quantified in two independent experiments.

### TBK1-ALS-associated mutations affect mitophagy through phosphorylation of TBK1 and OPTN

Several ALS-associated mutations and a number of engineered mutations have been shown to affect the structure, dimerization state, and kinase activity of TBK1 ^73^. Although mitochondrial recruitment of the TBK1 mutants during mitophagy has been assessed ^74^, the mechanisms underlying how ALS-associated mutations impact mitochondrial degradation remain to be elucidated. TBK1 dimerization is of great interest since the above findings indicate that assembly of the multimeric OPTN-TBK1 complex between the isolation membrane and damaged mitochondria is critical for TBK1 autophosphorylation. We thus examined the effects of various mutations on TBK1 function. R47H is an ALS-associated mutation of the kinase domain. D135N and S172A are engineered kinase inactive mutations, whereas G217R, R357Q, and M559R are ALS-associated mutations that impede dimerization ^73^. L693Q and V700Q are and engineered mutations that reduced *in vitro* interactions with ubiquitin ^59^. To monitor mutational effects on TBK1 stability, we used *TBK1-/-* HCT116 cells stably expressing YFP-P2A-TBK1 in which the P2A self-cleaving peptide was inserted between YFP and TBK1. Post-translation cleavage at this site would yield equivalent amounts of YFP and TBK1. Furthermore, we used a human codon-optimized TBK1 sequence for better expression (see materials and methods for details). Initially, cells expressing YFP-P2A-TBK1 WT and the various mutants were immunoblotted. As shown in Fig 5a, comparable YFP signals were detected among the YFP-P2A-TBK1 constructs, but the protein levels of the G217R, R357Q, and M559R mutants were reduced ∼ 50% compared to that of WT TBK1, suggesting that dimerization-deficient mutations destabilize TBK1. When inducing Parkin-mediated mitochondrial ubiquitination, OPTN in *TBK1-/-* cells and cells expressing either TBK1 WT or the mutants was efficiently ubiquitinated (Fig 5a), indicating that its association with damaged mitochondria in which Parkin is activated occurs regardless of TBK1. However, the levels of phosphorylated TBK1 (S172) and OPTN (S177) differed depending on the TBK1 mutant. L693Q and L700Q mutations induced TBK1 autophosphorylation comparable to WT TBK1, but the ALS-associated mutations R47H, G217R, and M559R were unable to phosphorylate either TBK1 or OPTN (Fig 5a, b). On the other hand, interestingly, while the protein level of TBK1 R357Q was 50% that of WT TBK1, phosphorylated TBK1 R357Q was also half (Fig 5a, b). Thus, monomeric TBK1 (especially R357Q) is unstable, but the mutant still has an ability to phosphorylate OPTN via TBK1 activation. We also noticed that the degree of TBK1 autophosphorylation correlated with OPTN phosphorylation at S177.

**Figure 5.**
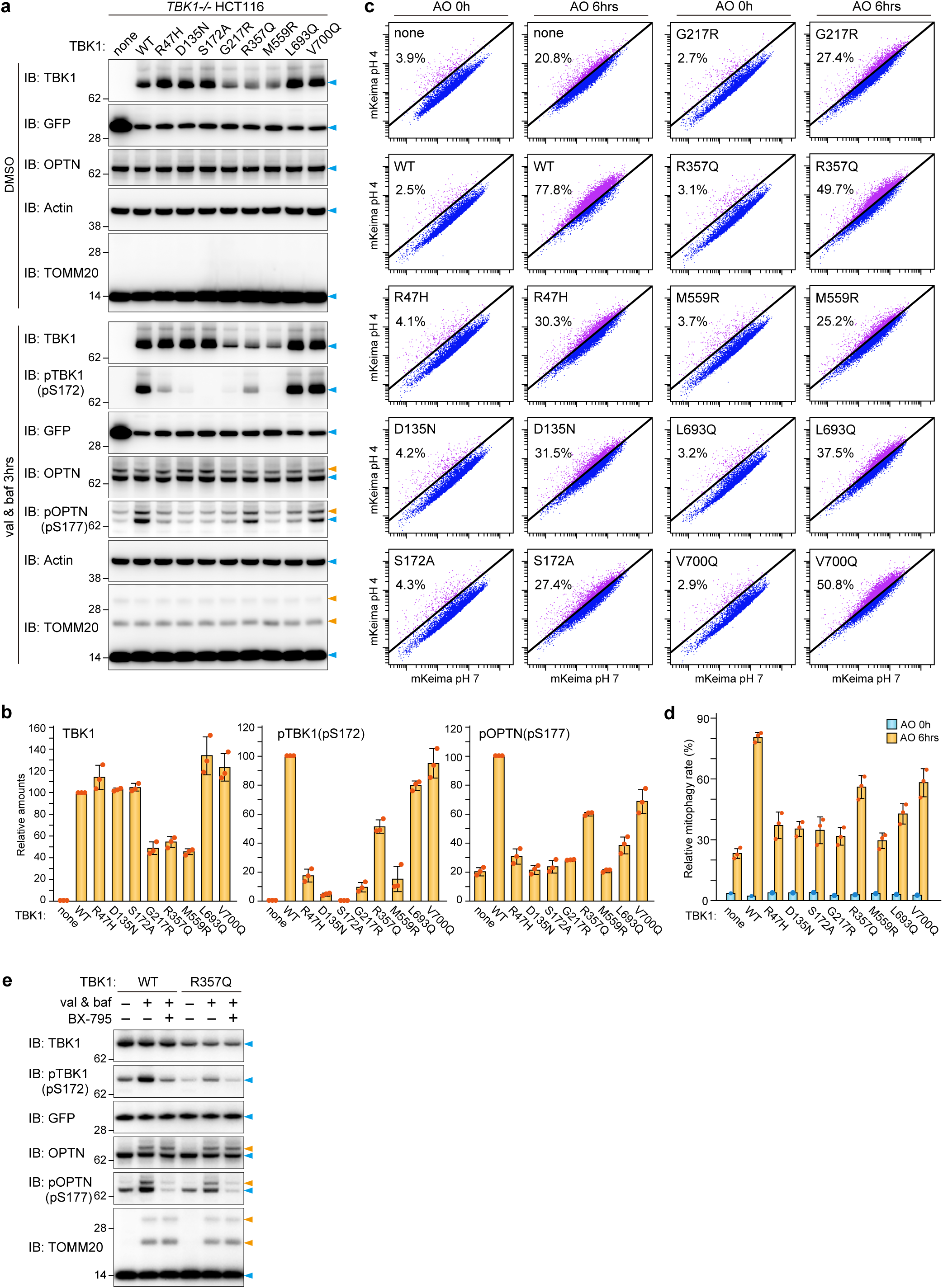
Impact of ALS-associated TBK1 mutations on mitophagy. **a** *TBK1-/-* HCT116 cells stably expressing Parkin, mt-Keima, and YFP alone (indicated as “none”) or YFP-P2A-TBK1 (WT or the indicated mutants) were treated with DMSO or valinomycin (val) and bafilomycin (baf) for 3 hrs and then analyzed by immunoblotting (IB). Orange arrowheads indicate ubiquitinated bands. **b** The levels of TBK1, pTBK1 and pOPTN after 3 hrs of val and baf treatment in (a) were quantified. The protein levels in cells expressing WT TBK1 were set to 100. Error bars represent mean ± s.d. of three independent experiments. **c** Cells in (a) were treated with antimycin A and oligomycin (AO) for 0 or 6 hrs and analyzed by FACS. Representative FACS data with the percentage of cells exhibiting lysosomal positive mt-Keima are shown. **d** Mitophagy rate (number of lysosomal positive Keima) in (c). Error bars represent mean ± s.d. of three independent experiments. **e** *TBK1-/-* HCT116 cells stably expressing Parkin and YFP-P2A-TBK1 (WT or the R357Q mutant) were treated with val and baf for 3 hrs with or without 2 μM BX-795. Total cell lysates were analyzed by IB. Orange arrowheads indicate ubiquitinated bands.

We next tested the effect of TBK1 mutations on mitochondrial degradation using Keima, a fluorescent protein with pH-dependent excitation spectra ^75^, as a reporter of mitophagy-induced delivery of mitochondria to the acidic lysosome lumen ^49, 56^. The fluorescence profile of cells expressing mitochondrial-targeted Keima was assessed via FACS after inducing mitophagy with oligomycin (AO) and antimycin A (Fig 5c). Mitophagy was evident in 78% of *TBK1-/-* HCT116 cells expressing YFP-P2A-TBK1 WT after 6 hrs with AO, whereas only 20% of *TBK1-/-* cells were mitophagy positive (Fig 5c). Although *in vitro* binding assays implicated L693Q and V700Q mutations in poly-Ub chain interactions ^59^, neither had a significant effect on mitophagy (Fig 5c, d). Similarly, TBK1 mutants (R47H, D135N, S172A, G217R, and M559R) that inhibited TBK1 and OPTN phosphorylation were unable to efficiently induce mitophagy (Fig 5c, d). Further, despite reduced protein levels of TBK1, the R357Q mutant could induce mitochondrial degradation (Fig 5c, d). In general, we found that the efficiency of mitophagy-induced mitochondrial degradation was correlated with the degree of S172 phosphorylation in TBK1.

There is a possibility that ULK1/2 may phosphorylate OPTN at S177 when TBK1 is mutated ^74^. To determine if this phosphorylation event is mediated by TBK1 R357Q or other kinases in response to mitophagy, we induced mitophagy in the presence of BX-795, a specific TBK1 kinase inhibitor. In *TBK1-/-* HCT116 cells expressing WT TBK1 in the presence of BX-795, the phosphorylation signals for OPTN S177 and TBK1 S172 were comparable to basal levels, suggesting complete inhibition (Fig 5e). Similar results were observed in *TBK1-/-* cells expressing the TBK1 R357Q mutant (Fig 5e). Therefore, TBK1 is the primary kinase for TBK1 autophosphorylation and OPTN phosphorylation during mitophagy even when TBK1 is monomeric.

### Creation of monobodies against OPTN

The results described above suggest that OPTN forms a contact site between isolation membranes and damaged mitochondria during Parkin-mediated mitophagy that supports TBK1 autophosphorylation. To determine if OPTN accumulation at the mitophagic contact site is required for TBK1 activation, we next sought to create monobodies that bind OPTN and physically prevent it from accumulating. Monobodies are non-immunogloblin-based proteins derived from the 10^th^ type III domain of human fibronectin that serve as backbone proteins that are sufficiently small for expression in both bacterial and mammalian cells ^76^. We previously developed a high-speed *in vitro* selection method termed TRAP (transcription-translation coupled with association of puromycin-linker) display for generating a monobody against a target protein ^77^. As shown in Fig 6a, monobody/mRNA complexes were synthesized from DNA pools in which the coding region for the BC-FG or CD-FG loops of a monobody were randomized. After reverse transcription, monobody/cDNA complexes were incubated biotinylated recombinant OPTN fragments and treated with streptavidin-coated magnetic beads, and the complexes with affinities for OPTN were selected. The amplified monobody DNAs were re-introduced into the TRAP system to yield a monobody library. We targeted two different N-terminal OPTN regions (26-196 aa and 133-196 aa), which is conjugated to biotin using an *in vitro* sortase reaction (Fig 6a and see materials and methods). After 7 or 8 round of selections, the recovered monobody cDNAs were sequenced (Supplementary Table 1). We selected 12 enriched clones from the pool for further analyses. YFP-fused monobodies were initially expressed in *NDP52* KO and *OPTN* KO HeLa cells, and a Keima-based mitophagy assay was performed. A previous study showed that both NDP52 and OPTN function as essential mitophagy adaptors and that either is sufficient for mitophagy ^49^, which means that *NDP52* KO or *OPTN* KO HeLa cells can drive Parkin-mediated mitophagy. Indeed, 79.3% of *NDP52* KO HeLa cells, in which OPTN is the lone mitophagy adaptor, were mitophagy-positive after 5 hrs with AO (Fig 6b, c). Interestingly, the expression of several OPTN monobody clones (clones #2, 3, 4, 5, 6, 8, 9) in *NDP52* KO cells inhibited mitophagy (Fig 6b, c). The inhibitory effects were specific to monobody-OPTN interactions as no mitophagy defects were observed in *OPTN* KO HeLa cells, which have NDP52 as the lone essential mitophagy adaptor (Fig 6d). To test if the monobodies can bind to OPTN in cells, *NDP52* KO HeLa cells stably expressing YFP alone or a YFP-monobody were co-immunoprecipitated. We confirmed that all of the YFP-monobodies tested were expressed (Fig 6e) and, with the exception of clones #1 and 11, that all interacted with endogenous OPTN (Fig 6f). Next, we characterized the OPTN monobodies *in vitro*. To verify the region of OPTN that interacts with the monobodies, the recombinant monobodies were pulled down using three different GST-OPTN constructs corresponding to OPTN aa 26-196, 26-119, or 133-196. All of the monobodies bound aa 26-196 of OPTN and all but clone #7 bound aa 26-119 (Fig 6g). The N-terminal region of OPTN (26-119 aa) interacts with the C-terminal region (677-729 aa) of TBK1 ^59^. GST-tagged TBK1 (677-729 aa) was incubated with EGFP or OPTN (26-196)-EGFP in the presence of the recombinant monobodies and the GST-TBK chimera was pulled down using glutathione sepharose (Fig 6h). For three clones (#3, 6, and 8), both OPTN (26-196)-EGFP and the monobodies were present in eluant, whereas no monobodies and reduced levels of OPTN (26-196)-EGFP were in the eluant for clones #4 and #9. These results indicate that the first group of clones (*i.e.,* #3, 6, and 8) can form a ternary complex with OPTN and TBK1 but that the other group of clones competitively prevent TBK1 from binding OPTN (Fig 6h, i).

**Figure 6.**
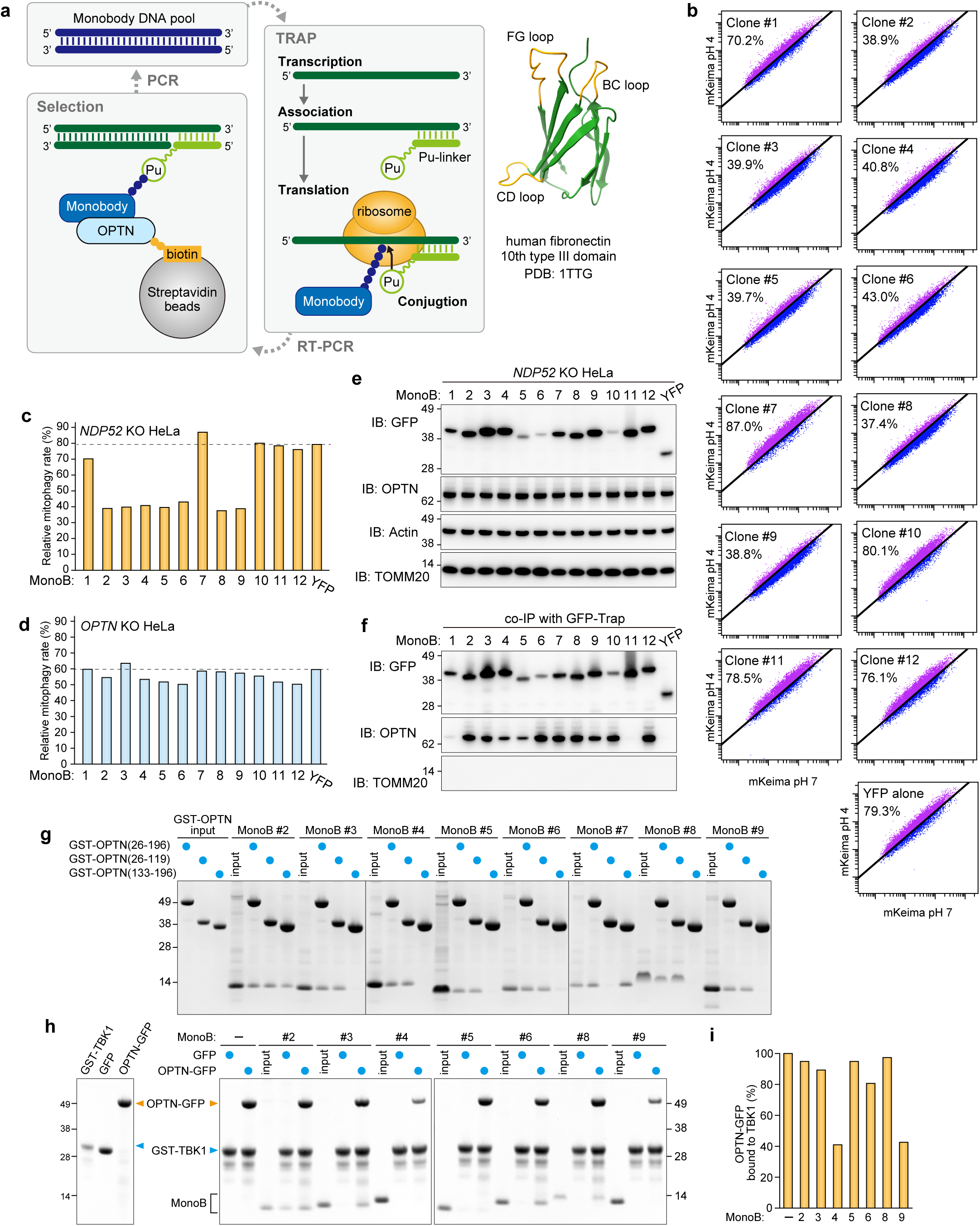
Selection and validation of OPTN monobodies. **a** Schematic representation of the process to select monobodies against OPTN using TRAP display. The BC, CD and FG loops in the monobody structure (PDB: 1TTG) are shown in orange. **b** *NDP52* KO HeLa cells expressing Parkin, mt-Keima, and YFP or YFP-tagged monobodies (MonoB) were treated with antimycin A and oligomycin (AO) for 5 hrs and analyzed by FACS. Representative FACS data with the percentage of cells exhibiting lysosomal positive mt-Keima are shown. **c** Mitophagy rate (number of lysosomal positive Keima) in (b). **d** Mitophagy rate (number of lysosomal positive Keima) using *OPTN* KO HeLa cells. **e** *NDP52* KO HeLa cells expressing Parkin and YFP or YFP-tagged MonoB were analyzed by immunoblotting (IB). **f** The cells in (e) were co-immunoprecipitated (co-IP) using GFP-Trap. The bound fractions were analyzed by IB. **g** Recombinant GST-OPTN (26-196aa, 26-119aa, or 133-196aa) were incubated with recombinant MonoB. The glutathione sepharose bound fractions were analyzed by CBB staining. **h** Recombinant GST-TBK1 (677-729aa) and MonoB were incubated with of recombinant GFP or OPTN (26-196aa)-GFP. GST-TBK1 (677-729aa) was pulled down with glutathione sepharose and the bound fractions were analyzed by CBB staining. **i** The levels of OPTN (26-196aa)-GFP pulled down with GST-TBK1 (677-729aa) in (h) were quantified. The level of OPTN (26-196aa)-GFP pulled down without MonoB was set to 100.

### Monobodies against OPTN inhibit both the assembly and accumulation of OPTN at the mitophagic contact site

We selected monobody clones #3 and #4 for further analysis. First, the binding constants for OPTN and the monobodies were assessed using biolayer interferometry (BLI). OPTN (26-196 aa) was immobilized on the streptavidin-coated sensor and various concentrations of the monobodies were applied to determine the kinetic parameters based on global fitting (Supplementary Fig 3). Both clones exhibited nanomolar affinities (clone #3, Kd = 4.56 nM; clone #4, Kd = 6.19 nM) for the OPTN N terminus, which are much lower (*i.e.,* stronger) than the 2.5 μM reported for TBK1 (677-729 aa) and OPTN (26-119 aa) ^59^. Next, we examined how the monobodies block mitophagy. For these assays, the two monobodies were stably expressed in *NDP52* KO HeLa cells and the cellular localization of endogenous OPTN during mitophagy was determined (Fig 7a). As a control, we also examined the localization of OPTN in *NDP52* KO cells expressing GFP. In these cells, OPTN formed dots and sphere-like structures on damaged mitochondria. In sharp contrast, this OPTN distribution was not observed in cells expressing the GFP-tagged monobodies, however, mitochondrial recruitment of OPTN was still observed during mitophagy (Fig 7a). Furthermore, immunoblots showed that expression of the monobodies abolished both autophosphorylation of S172 in TBK1 and S177 phosphorylation in OPTN (Fig 7b, c). Mitophagy-induced ubiquitination of OPTN was not disrupted by the presence or absence of the monobodies (Fig 7b), which is consistent with mitochondrial recruitment of OPTN regardless of monobody expression.

**Figure 7.**
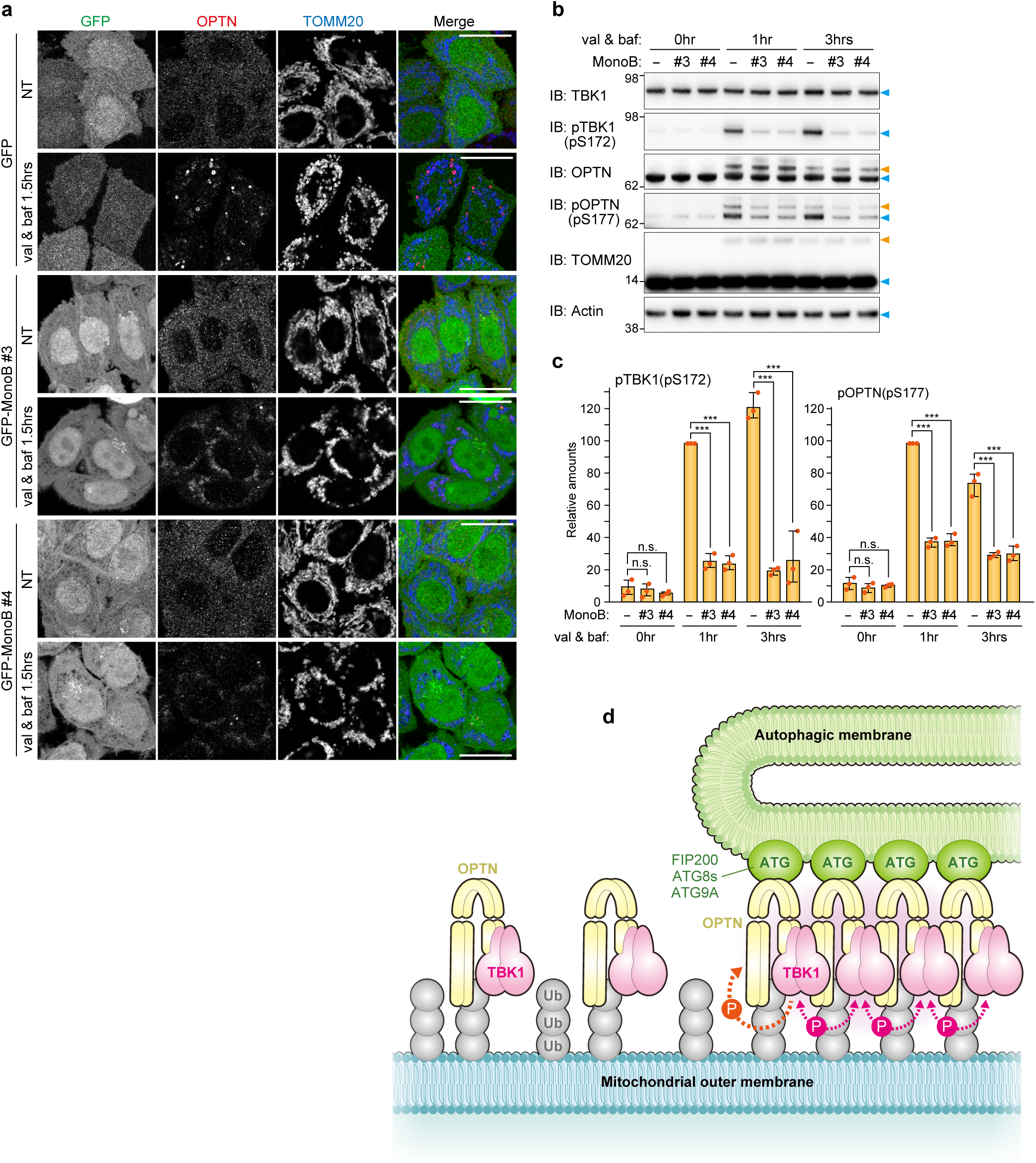
OPTN monobodies inhibit OPTN assembly at the mitophagy-contact sites. **a** *NDP52* KO HeLa cells expressing Parkin, GFP alone, or GFP-tagged monobodies (MonoB) were treated with or without valinomycin (val) and bafilomycin (baf) for 1.5 hrs. The cells were immunostained with anti-OPTN and TOMM20 antibodies. Bars, 10 μm. **b** The cells in (a) were analyzed by immunoblotting (IB). Orange arrowheads indicate ubiquitinated bands. **c** The levels of pTBK1 and pOPTN in (b) were quantified. The protein levels in cells without MonoB after 1 hr val and baf treatment were set to 100. Error bars represent mean ± s.d. of three independent experiments. **d** Proposed model of TBK1 activation during PINK1/Parkin-mediated mitophagy.

## Discussion

TBK1 is a key regulatory kinase in the cell signaling pathways underlying innate immune responses and with elimination of invader pathogens and damaged mitochondria ^78^. To modulate this signaling, TBK1 can bind to different adaptor proteins. These interactions facilitate local clustering of TBK1 and allow the interdimer kinase domains of TBK1 to activate via trans-autophosphorylation ^79^. TBK1 is a multimeric domain protein consisting of a kinase domain, a ubiquitin-like domain, a scaffold dimerization domain, and C-terminal domain that can bind various adaptor proteins. TBK1 forms a rigid dimer in which the two kinase catalytic sites are oriented away from one another. Thus, TBK1 autophosphorylation requires at least dimerization of homo TBK dimer. In the STING pathway, the binding of cGAMP by STING induces the oligomerization of STING on post-Golgi membranes ^80, 81^. Subsequent higher-order STING oligomers arranged in a linear manner and a C-terminal STING segment bind TBK1 dimers without structural changes in TBK1. Therefore, STING oligomers provide a platform for TBK1 oligomerization, which induces trans-autophosphorylation of adjacent TBK1 proteins. Thus, in the STING pathway, STING oligomerization directly induces TBK1 activation.

During Parkin-mediated mitophagy, OPTN recruits TBK1 onto damaged mitochondria in a Parkin-dependent manner, which had been thought to induce TBK1 activation ^62, 63^. However, we showed that OPTN recruitment to damaged mitochondria, itself, is insufficient for TBK1 autophosphorylation. Although OPTN forms a dimer through the coiled-coil domains, OPTN cannot oligomerize on its own even on ubiquitin-decorated mitochondria. In this study, we showed that TBK1 autophosphorylation requires OPTN for interactions with both the autophagy machinery and the ubiquitin-coated cargo. Previously, it was thought that TBK1 activates via autophosphorylation before phosphorylating OPTN to recruit autophagic membranes to mitochondria. However, our results show that formation of the isolation membrane itself is involved in TBK1 autophosphorylation. We show that OPTN interacts with both autophagy machinery and ubiquitin to form contact sites that can accumulate TBK1. Further, our results show that autophagic membrane formation affects TBK1 autophosphorylation, which must occur prior to TBK1 activation, thereby negating the model in which TBK1 activation results in the phosphorylation of OPTN S177 and the subsequent recruitment of autophagic membranes to mitochondria. Signaling cascades for most kinases are unidirectional (*i.e.,* the transfer of signals from upstream components to those downstream). In response to PINK1/Parkin-mediated mitophagy, the initial TBK1 autophosphorylation step requires downstream interactions between OPTN and the autophagy components, suggesting a positive feedback loop. TBK1 phosphorylation of the OPTN UBAN domain (*e.g.* S473) promotes tight association of OPTN with ubiquitin-coated mitochondria, whereas TBK1 phosphorylation of S177 in the LIR motif induces translocation of mitochondria-associated OPTN to the autophagosome formation site. This accumulation of OPTN at a contact site between the autophagy machinery and ubiquitin-coated mitochondria provides a platform for TBK1 hetero-autophosphorylation by adjacent TBK1 at the contact site (Fig 7d). Activated TBK1 then phosphorylate other OPTN-TBK1 pairs, which are newly recruited to damaged mitochondria, and the next phosphorylation cycle proceeds. Thus, OPTN-mediated TBK1 phosphorylation and TBK1-mediated OPTN phosphorylation appear to form a self-propagating positive feedback loop, that leads to elongation of isolation membranes on damaged mitochondria. Intervention of OPTN monobodies also support this model. Although monobody clone #3 formed a ternary complex with OPTN and TBK1 (Fig 6h), it prevented OPTN from accumulating, which abolished formation of the mitophagic contact site and subsequent TBK1 autophosphorylation.

Some of our OPTN monobodies (such as clones #4 and #9) directly disturbed OPTN-TBK1 interactions. Interestingly, an OPTN E50K mutation was identified in familial primary open-angle glaucoma ^82^ that promotes stronger interactions with TBK1 ^59^, which in turn triggers an accumulation of insoluble OPTN and constriction of the Golgi body ^83^. Our monobodies could potentially be used to suppress the disease-related phenotype by weakening the interaction between OPTN and TBK1. This hypothesis, however, will need to be tested in for future studies.

TBK1 mediates phosphorylation of various Ser/Thr sites on at least four autophagy adaptors, OPTN, NDP52, TAX1BP1, and p62 ^63^. Although phosphorylated S177 and S473 in OPTN enhance interactions with LC3 and K63-linked ubiquitin chains, respectively, other sites, including S262, appear to be randomly phosphorylated. Furthermore, p62, which is recruited to mitochondria but does not accumulate at the mitophagic contact site (Supplementary Fig 1), was also highly phosphorylated by TBK1 during mitophagy. During manuscript preparation, Nguyen *et al.* reported direct interactions between TBK1 and the PI3K complex facilitate mitophagy initiation by OPTN (doi: https://doi.org/10.1101/2022.08.14.503930). Our results support this finding as recruitment of OPTN, but not NDP52, to the autophagosome formation site strictly depends on TBK1 (Fig 1e, f). Furthermore, that study showed that OPTN-mediated mitophagy does not require the ULK1/2 kinases and that TBK1 compensates for ULK kinases. Based on our findings and those of other researchers, TBK1 does not appear to have strict substrate specificity requirements. Rather, the most critical aspect of TBK1 is the spatial location of activation (*i.e.,* at the autophagosome formation site on mitochondria). Since OPTN-TBK1 localizes to the mitophagic contact site where autophagy core components also accumulate, it seems to follow that activated TBK1 would be able to non-discriminately phosphorylate autophagy core components in close proximity.

### Materials and methods

Reagents including cell lines, antibodies, and plasmid DNAs including siRNA used in this study are listed in Tables below.

### Plasmid Construction

Human codon-optimized *TBK1* was synthesized by eurofins genomics and was inserted into the EcoRI site of pMXs-puro_EYFP-P2A-EcoRI to make pMXs-puro_YFP-P2A-hOpt-TBK1. Mutations were introduced by PCR-based DNA mutagenesis.

### Cell culture and transfection

HeLa and HEK293T cells were cultured in DMEM supplemented with 10% (v/v) FBS, 1 mM sodium pyruvate, nonessential amino acids, and PSG. HCT116 cells were cultured in McCoy’s 5A medium supplemented with 10% (v/v) FBS, nonessential amino acids, and 2 mM GlutaMax. The Penta KO HeLa ^49^, *NDP52* KO HeLa ^49^, *OPTN* KO HeLa ^49^, *FIP200* KO HeLa ^55^, *ATG9A* KO HeLa ^84^, *ATG5* KO HeLa ^84^, and *TBK1-/-* HCT116 ^56^ cell lines were described previously.

The HeLa AAVS-Parkin cell line was established as follows. A donor Parkin plasmid was constructed using a pZDonor-AAVS Puromycin Vector Kit. The resultant donor plasmid was transfected with AAVS TALEN plasmids into HeLa cells using FuGENE6. The cells were grown in the presence of 1 μg/ml puromycin and single clones were isolated into 48-well plates. Insertion of the *PARKIN* gene into the AAVS locus was confirmed by isolation of genomic DNA followed by PCR. Parkin expression and mitochondrial recruitment were confirmed by immunostaining valinomycin-treated cells with an anti-Parkin antibody.

Stable cell lines were made by recombinant retrovirus infection ^56^. FuGENE6 and FuGENE HD reagents were used for plasmid transfection according to the manufacturer’s instructions. RNAiMAX was used for siRNA transfection.

Final concentrations of reagents used were: 10 μM valinomycin, 10 μM oligomycin, 4 μM antimycin A, 100 nM bafilomycin A1, 1 μM Epoxomicin, 10 μM MG132 and 100 nM Concanamycin A. To block apoptotic cell death in response to Parkin-mediated ubiquitination, 5μM Q-VD-OPH was added.

### CRISPR/Cas9-edited gene knockout

*TBK1* KO, *FIP200/TBK1* DKO, and *ATG9A/TBK1* DKO HeLa cell lines were established via CRISPR/Cas9-based genome editing. The three gRNA target sequences (5’-GAC CCT TTG AAG GGC CTC GTA GG-3’, 5’-ATT CCT ACG AGG CCC TTC AAA GG -3’ and 5’-GGC CCT TCA AAG GGT CTA AAT GG -3’) for *TBK1* exon 6 were designed using CRISPOR (http://crispor.tefor.net/). Each of the oligonucleotide pairs were annealed and introduced into the BpiI site of the PX459 vector to yield PX459-TBK1-ex6-1, PX459-TBK1-ex6-2, and PX459-TBK1-ex6-3. The plasmids were transfected into WT HeLa, *FIP200* KO HeLa, and *ATG9A* KO HeLa cell lines. Puromycin-resistant cells were seeded into 96 well plates, and single clones were analyzed by immunoblotting to confirm *TBK1* knockout.

### Immunoblotting

Cells grown in 6-well plates were washed twice with Phosphate Buffered Saline (PBS) and solubilized with 2% CHAPS buffer (25 mM HEPES-KOH pH7.5, 300 mM NaCl, 2% [w/v] CHAPS, cOmplete) on ice for 10 min. To detect phosphorylated proteins, PhosSTOP was added to the 2% CHAPS buffer. After centrifugation at 12,000 ×g for 2 min at 4°C, the supernatants were collected, and protein concentrations were determined on a DS-11+ spectrophotometer (DeNovix). SDS-PAGE sample buffer with DTT was added to the supernatants, which were then incubated at 42°C for 5 min. To detect hAG constructs, the samples were boiled at 95°C for 5 min. Total cell lysates were loaded on NuPAGE 4-12% Bis-Tris gels and electrophoresed using MES running buffer. Proteins were transferred to PVDF membranes that were blocked with 2% (w/v) skim-milk/TBS-T and then incubated with primary and HRP-conjugated secondary antibodies. Proteins were detected using a Western Lighting Plus-ECL Kit on an ImageQuant LAS4000 (GE Healthcare) or a FUSION SOLO S system (VILBER). ImageJ was used to quantify protein bands.

### Immunostaining and immunofluorescence microscopy

Cells grown on glass bottom 35-mm dishes (MatTek) were fixed with 4% PFA solution for 20 min at room temperature, permeabilized with 0.15% (v/v) TritonX-100 in PBS for 20 min, and preincubated with 0.1% (w/v) gelatin in PBS for 30 min. The cells were incubated with primary antibodies diluted in 0.1% (w/v) gelatin for 2 hrs and Alexa Fluor-conjugated secondary antibodies diluted in 0.1% (w/v) gelatin for 1 hr. Microscopy images were captured using an inverted confocal microscope (LSM710 and LSM780, CarlZeiss) with a Plan-Apochromat 63x/1.4 Oil DIC lens. For image analysis, ZEN microscope software and Photoshop (Adobe) were used.

### Phos-tag PAGE

Phos-tag PAGE analysis was described previously ^42^.

### Selection of monobodies against OPTN

*Escherichia coli* codon-optimized *OPTN* DNA encoding aa 26-196, 26-119, or 133-196 was synthesized by eurofins genomics and then inserted into the BamHI site of pGEX6P1 together with the sortase recognition sequence and a His6-tag sequence to generate PGEX6P1_Opt-OPTN (26-196 aa, 26-119 aa, or 133-196 aa)-Sortase-His6-STOP. The plasmids were introduced into *E. coli* BL21-CodonPlus(DE3)-RIL competent cells, and the transformants were grown in LB medium supplemented with 100 μg/ml ampicillin and 25 μg/ml chloramphenicol at 37°C. OPTN-sortase-His6 was expressed at 18°C overnight by adding 200 μM IPTG. Bacterial cell pellets were resuspended in TBS buffer (50 mM Tris-HCl pH7.5, 120 mM NaCl) supplemented with lysozyme, DNAse I, DTT, MgCl_2_, and cOmplete and stored at -20°C. Frozen cell suspensions were thawed and sonicated (Advanced-Digital Sonifer, Branson) and insoluble proteins removed by centrifugation. The supernatants were mixed with equilibrated glutathione sepharose for 40 min at 4°C and then loaded onto a column and washed with TBS buffer. The sepharose columns were incubated overnight at 4°C with Prescission protease. GST-cleaved OPTN-sortase-His proteins were subsequently collected. Biotin-modified OPTN was prepared as follows. OPTN-sortase-His was incubated with NH_2_-GGG(Lys[Biotin])-CONH_2_ (eurofins) and His-tagged Sortase 5Y (a kind gift from Dr. Yasushi Saeki) in TBSC buffer (50 mM Tris-HCl pH7.5, 120 mM NaCl, 5mM CaCl_2_) for 1.5 hrs at 37°C. Unmodified OPTN-sortase-His and His-tagged Sortase 5Y were removed by TALON resin and GGGK-biotin was removed using PD Miditrap G-25.

The monobody mRNA libraries were prepared using similar procedure as reported before ^77^ and the detailed procedure will be reported elsewhere. For the first round of selection, 1 μM mRNA library/HEX-mPuL was added to a reconstituted translation system, and the reaction mixture (500 μL) was incubated at 37°C for 10 min. After the reaction, 41.7 μL of 200 mM EDTA (pH 8.0) was added to the translation mixture. Reverse transcription mixture (144.3 μL; 224 mM Tris–HCl, pH 8.4, 336 mM KCl, 96 mM MgCl_2_, 22 mM DTT, 2.3 mM each dNTP, 6.9 μM MoS-G5R-T.R28 [5’-ACT ATC GGC CTC CTC CTC CAC CTT GAC T-3’] primer, and 5.5 μM MMLV) was added to the translation mixture, and the resulting solution was incubated at 42°C for 15 min. The buffer was changed to HBST buffer (50 mM Hepes-KOH pH 7.5, 300 mM NaCl, and 0.05% [v/v] Tween 20) using Zeba Spin Desalting Columns. To remove the bead binders, the resulting solution was mixed with 100 μL of Dynabeads M-280 streptavidin (Thermo Fisher Scientific) at 25°C for 10 min. The supernatant was mixed with 28.8 μL of target protein mixture containing 0.5 μM each biotinylated OPTN fragment at a final concentration of 20 nM. The resulting solution was incubated at 25°C for 10 min. The target proteins were collected by mixing with 41.1 μL of Dynabeads M-280 streptavidin for 1 min. The collected beads were washed with 500 μL of the HBST buffer for 1 min two times, and 690 μL of PCR premix (10 mM Tris–HCl pH 8.4, 50 mM KCl, 0.1% [v/v] Triton X-100, 2 mM MgCl_2_, and 0.25 mM each dNTP) was added. The beads were heated at 95°C for 5 min, and the amount of eluted cDNA was quantified by SYBR green-based quantitative PCR using T7SD8M2.F44 (5’-ATA CTA ATA CGA CTC ACT ATA GGA TTA AGG AGG TGA TAT TTA TG-3’) and MoS-RealTime.R20 (5’-AGC ATC CCA GCT GAT CTG AA-3’) as primers. The eluted cDNA was PCR-amplified using T7SD8M2P.F47 (5’-ATA CTA ATA CGA CTC ACT ATA GGA TTA AGG AGG TGA TAT TTA TGC CT-3’), Mos-G5R-T-an21-3.R45 (5’-CCC GCC TCG CGC CCG CCG TCC ACT ATC GGC CTC CTC CTC CAC CTT-3’) and Pfu-S DNA polymerase and purified by phenol/chloroform extraction and isopropanol precipitation. From the following selection, the resulting DNA (1.25 nM final concentration) was added to the TRAP system, and the reaction mixture (10 μL) was incubated at 37°C for 30 min. The other procedure was similar to the above description. After the final round of selection, the sequences of the recovered DNA were analyzed using an Ion Torrent instrument (Thermo Fisher Scientific). DNAs encoding the monobodies were synthesized (eurofin genetics) and subcloned into pMXs-puro_YFP-TEV, pMXs-puro_GFP-TEV, and pET21a(+) vectors.

### Mitophagy assay using mito-Keima and FACS

FACS-based mitophagy measurements were performed as follows. For *TBK1-/-* HCT116 cells, untagged Parkin, YFP-P2A-TBK1, and mt-Keima were stably expressed via retrovirus infections. For *OPTN* KO and *NDP52* KO HeLa cells, untagged Parkin, YFP-monobody, and mt-Keima were stably expressed. Cells were grown in a 6-well plate and treated with 4 μM antimycin A and 10 μM oligomycin (AO) for varying lengths of time. Trypsinized cells were resuspended in PBS containing 2.5% FBS. FACS analysis was performed using a BD LSRFortessa X-20 cell sorter (BD Biosciences) with FACSDiva software. Keima fluorescence was measured as a dual-excitation ratiometric pH system using 405-nm (pH 7) and 561-nm (pH 4) lasers and 610/20-nm emission filters. Each sample measured consisted of 10,000 YFP/Keima double positive cells.

### Preparation of recombinant proteins

Recombinant OPTN monobodies, OPTN (26-196aa)-EGFP, EGFP, and GST-TBK1 (677-729aa) were obtained as follows. pET21a(+)_OPTN MonoS, pET16b_opt-OPTN (26-196aa)-EGFP, pET16b_EGFP and pGEX6P1_opt-TBK1 (677-729aa) were introduced into *E. coli* BL21-CodonPlus(DE3)-RIL competent cells, and the transformants were grown in LB medium supplemented with 100 μg/ml ampicillin and 25 μg/ml chloramphenicol at 37°C. The E50K mutation was introduced into OPTN (26-196aa) to enhance interactions with TBK1 ^59^. His6-tagged monobody, His10-tagged OPTN (26-196aa)-EGFP, His10-tagged EGFP, and GST-TBK1 (677-729aa) were expressed overnight at 20°C with 100 μM IPTG. Bacterial cell pellets were resuspended in TBS buffer with lysozyme, DNAse I, DTT, MgCl_2_, and cOmplete and stored at -20°C until used. The frozen cell suspension was thawed, sonicated, and insoluble proteins were removed by centrifugation. His-tagged protein supernatants were mixed with equilibrated Ni-NTA, whereas GST-tagged proteins supernatants were mixed with equilibrated glutathione Sepharose. Protein supernatant mixtures were incubated for 30 min at 4°C and then washed with TBS buffer. His-tagged proteins were eluted stepwise with imidazole. GST-tagged proteins were eluted with 20 mM glutathione. The elution buffers were exchanged for 50 mM Tris-HCl pH7.5, 120 mM NaCl, 1 mM TCEP, 10%(w/v) glycerol, and the proteins were stored at -80°C.

### Pull down assay using recombinant proteins

Recombinant proteins GST-TBK1 (677-729 aa), OPTN (26-196 aa)-EGFP, EGFP alone, and monobodies were incubated as 4.5 μM aliquots with equilibrated glutathione Sepharose in 100 μL binding buffer (50 mM Tris-HCl pH7.5, 120 mM NaCl, 1mM TCEP, 10%[w/v] glycerol) for 30 min at room temperature. The resin was washed three times with 1 mL of binding buffer containing 0.1% (v/v) TritonX-100 and bound proteins were eluted with SDS-PAGE sample buffer.

### Kd determination

Protein affinities were determined for biotinylated OPTN (26-196 aa) immobilized on a streptavidin biosensor (ForteBio) using the Octet system (ForteBio) according to the manufacturer’s instructions. The analyte monobody was initially dissolved in 50 mM Tris-HCl pH7.5, 120 mM NaCl, 1 mM TCEP, and 200 mM imidazole and then was exchanged to buffer D (50 mM Hepes-KOH, pH 7.5, 300 mM NaCl, 0.05% [v/v] Tween 20, and 0.1% [w/v] PEG6000) using Zeba Spin Desalting Columns. The protein concentration was measured via A280 according to the molar extinction coefficient estimated from the amino acid composition. The binding assay was performed at 30 °C in buffer D. The binding assay steps consisted of equilibration for 150 s, association for 600 s, and dissociation for 600 s.

### Coimmunoprecipitation using GFP-Trap

*NDP52* KO HeLa stably expressing YFP alone or YFP-tagged monobodies grown on 35-mm dishes were washed with PBS and solubilized with 0.2% TX-100 buffer (50mM Tris-HCl pH7.5, 150 mM NaCl, 0.2%(v/v) TritonX-100, cOmplete) on ice for 20 min. After centrifugation (16,000 ×g for 10 min at 4°C), the supernatants were incubated with equilibrated GFP-Trap agarose for 1 hr at 4°C. The agarose was three times washed with 0.2% TX-100 buffer and the proteins were eluted with SDS-PAGE sample buffer.

### Statistical analysis

Statistical analysis was performed using data obtained from three or more biologically independent experimental replicates. A t-test was used for statistical comparisons between two groups (n.s., not significant; *, P < 0.05; **, P < 0.01; ***, P < 0.001).

Table for Reagents and cell lines

| REAGENT or RESOURCE | SOURCE | IDENTIFIER |
| --- | --- | --- |
| Reagents |  |  |
| Antimycin A | Sigma-Aldrich | Cat. # A8674 |
| Oligomycin | Calbiochem | Cat. # 495455-10MGCN |
| Valinomycin | Sigma-Aldrich | Cat. # V0627 |
| Bafilomycin A1 | Calbiochem | Cat. # 196000 |
| DMEM | Sigma-Aldrich | Cat. # D5796 |
| McCoy's 5A | Gibco | Cat. # 16600-082 |
| FBS | biowest | Cat. # S1820 |
| MEM Non-Essential Amino Acids Solution | Gibco | Cat. # 11140-050 |
| Sodium Pyruvate | Gibco | Cat. # 11360-070 |
| Penicilin-Streptomycin-Glutamine | Gibco | Cat. # 10378-016 |
| GlutaMAX Supplement | Gibco | Cat. # 35050-061 |
| Trypsin-EDTA (0.25%) | Gibco | Cat. # 25200-056 |
| FuGENE6 | Promega | Cat. # E269A |
| Lipofectamine LTX | Invitrogen | Cat. # 15338100 |
| Lipofectamine RNAiMAX | Invitrogen | Cat. # 13778150 |
| CHAPS | Sigma-Aldrich | Cat. # C3023 |
| cOmplete, EDTA-free Protease Inhibitor Cocktail | Roche | Cat. # 11873580001 |
| PhosSTOP, Phosphatase Inhibitor Cocktail | Roche | Cat. # 04906845001 |
| BX-795 (TBK1 inhibitor) | abcam | Cat. # ab142016 |
| 4% PFA Solution | Wako | Cat. # 163-20145 |
| Gelatin | Sigma-Aldrich | Cat. # G9391 |
| Triton X-100 | MP Biomedicals | Cat. # 807426 |
| Tween 20 | nacalai tesque | Cat. # 35624-15 |
| NuPAGE 4-12% Bis-Tris gel | Invitrogen | Cat. # NP0323BOX |
| Difco Skim Milk | BD Biosciences | Cat. # 232100 |
| Phusion High-Fidelity DNA polymerase | Thermo Fisher Scientific | Cat. # F-530L |
| DNA Ligation Kit | TaKaRa | Cat. # 6023 |
| Western Lightning Plus-ECL | PerkinElmer | Cat. # NEL105001EA |
| ECL Prime Western Blotting Detection Reagents | GE Healthcare | Cat. # RPN2232 |
| BL21-CodonPlus(DE3)-RIL Competent Cells | Agilent Technology | Cat. # 230245 |
| FuGENE® HD Transfection Reagent | Promega | Cat. # E2311 |
| TCEP (Tris(2-carboxyethyl)phosphine hydrochloride) | Sigma | Cat. # C4706-10G |
| Ni-NTA Agarose | QIAGEN | Cat. # 30230 |
| Glutathione Sepharose 4B | cytiva | Cat. # 17075605 |
| PD MidiTrap™ G-25 | cytiva | Cat. # 28918008 |
| PreScission™ Protease | cytiva | Cat. # 27084301 |
| DTT (1,4-dithiothreitol) | Roche | Cat. # 10708984001 |
| Imidazole | FUJIFILM | Cat. # 095-00015 |
| L-Glutathione reduced | Sigma | Cat. # G4251-25G |
| GFP-Trap Agarose | ChromoTek GmbH | Cat. # gta-20 |
| Epoxomicin | PEPTIDE INSTITUTE, INC. | Cat. # 4381-v |
| Z-Leu-Leu-Leu-H (aldehyde) MG-132 | PEPTIDE INSTITUTE, INC. | Cat. # 3175-v |
| Concanamycin A | Adipogen Life Sciences | Cat. # BVT-0237-C100 |
| Cell lines |  |  |
| WT HeLa | ATCC | CCL-2,2 |
| HeLa AAVS-Parkin | This study |  |
| *FIP200* KO HeLa | (Vargas et al., 2019) |  |
| *ATG5* KO HeLa | (Nezich et al., 2015) |  |
| *ATG9A* KO HeLa | (Nezich et al., 2015) |  |
| Penta (OPTN/NDP52/TAX1BP1/p62/NBR1) KO HeLa | (Lazarou et al., 2015) |  |
| *TBK1* KO HeLa | This study |  |
| *FIP200/TBK1* DKO HeLa | This study |  |
| *ATG9A/TBK1* DKO HeLa | This study |  |
| HEK293T | ATCC | CRL-3216 |
| HCT116 | (Yamano et al., 2018) |  |
| *TBK1*<sup>-/-</sup> HCT116 | (Yamano et al., 2020) |  |

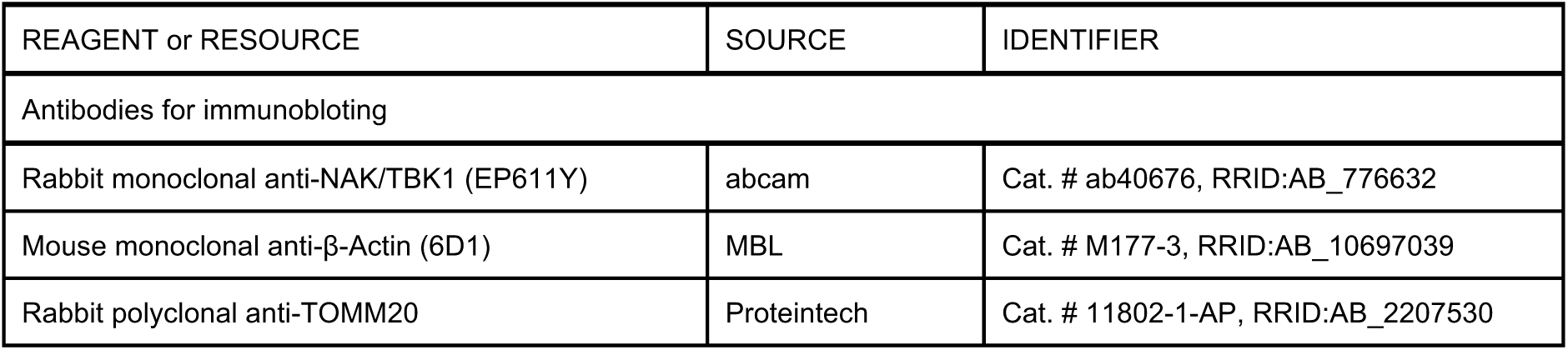
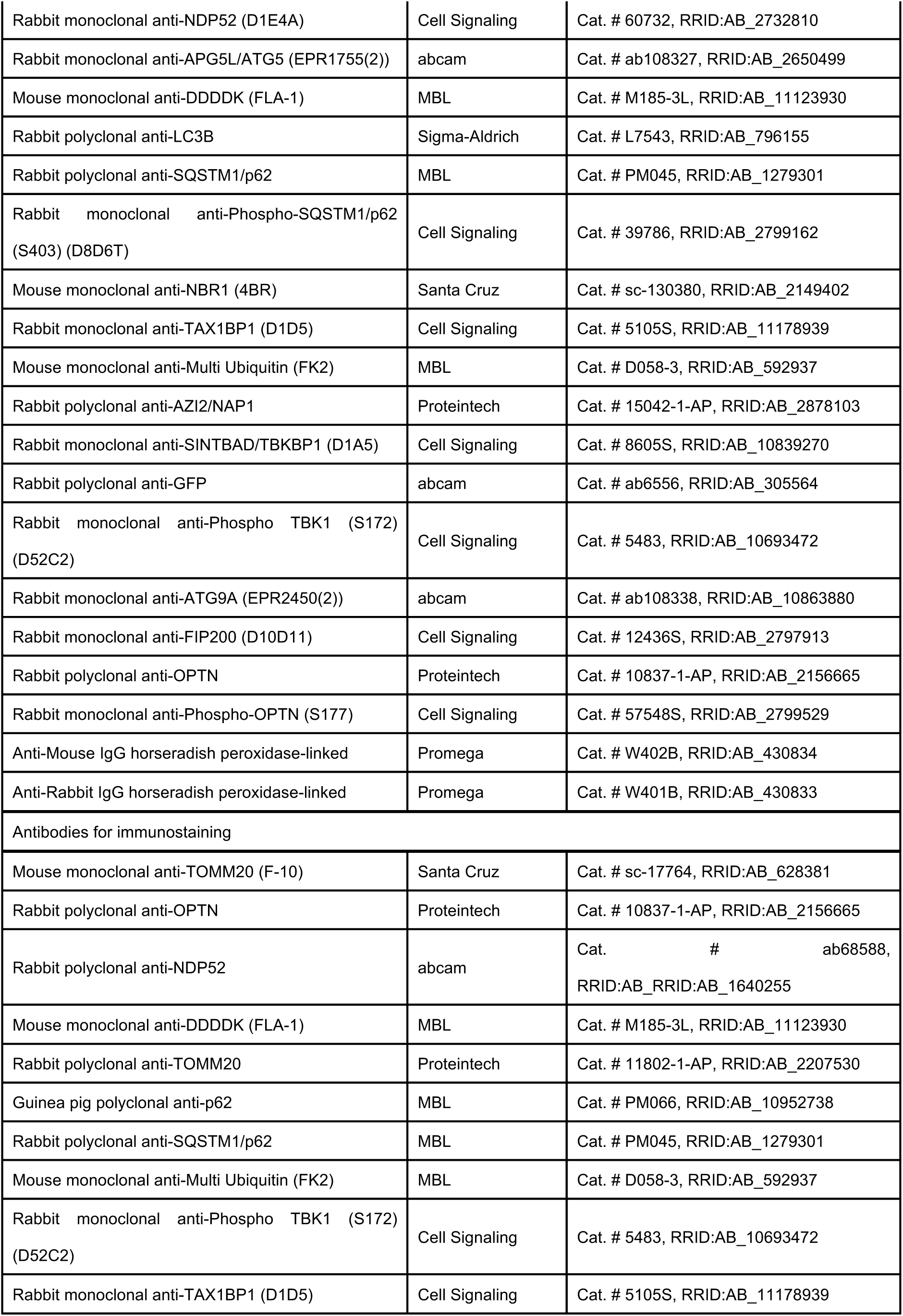
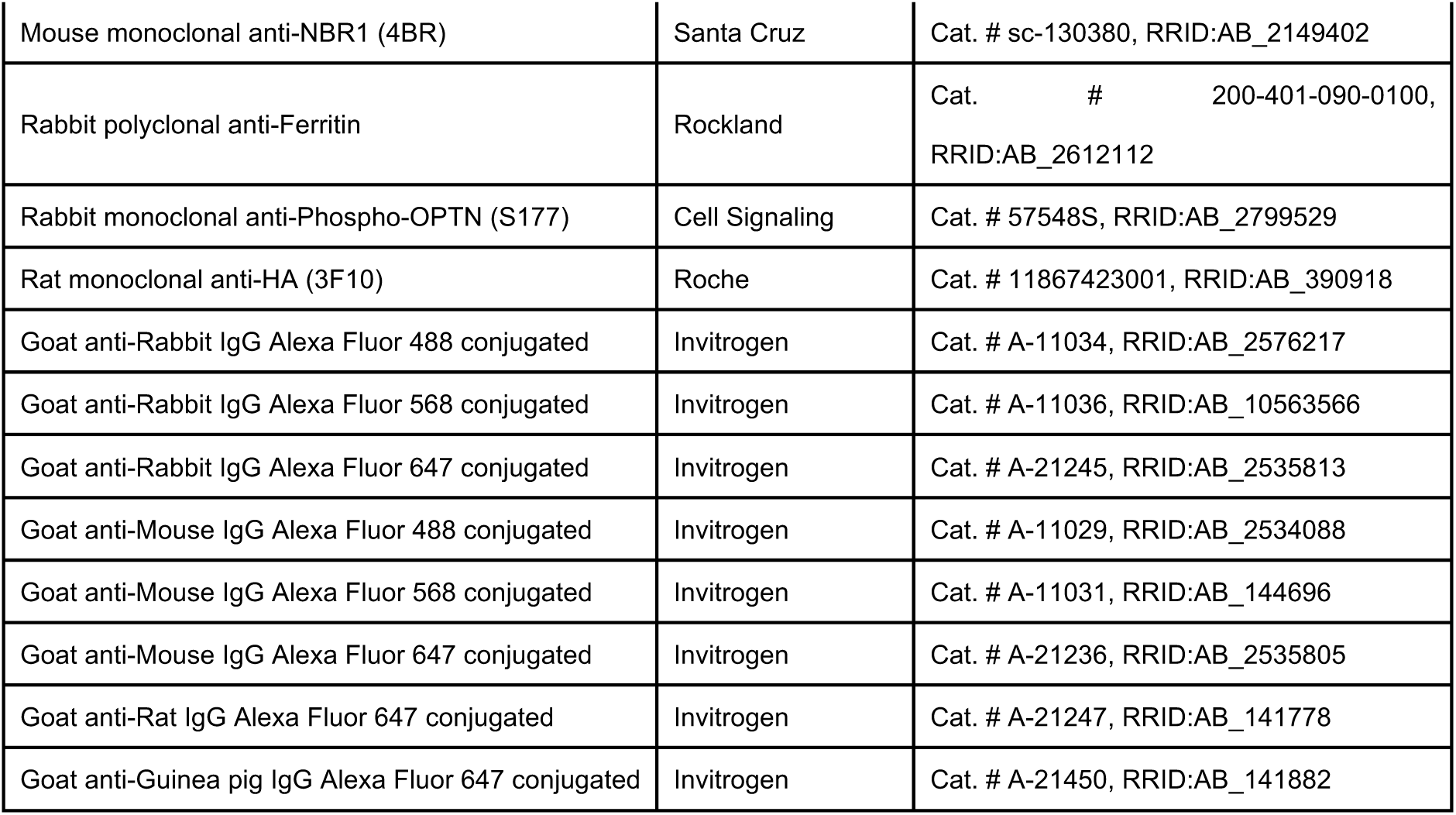
Table for Antibodies

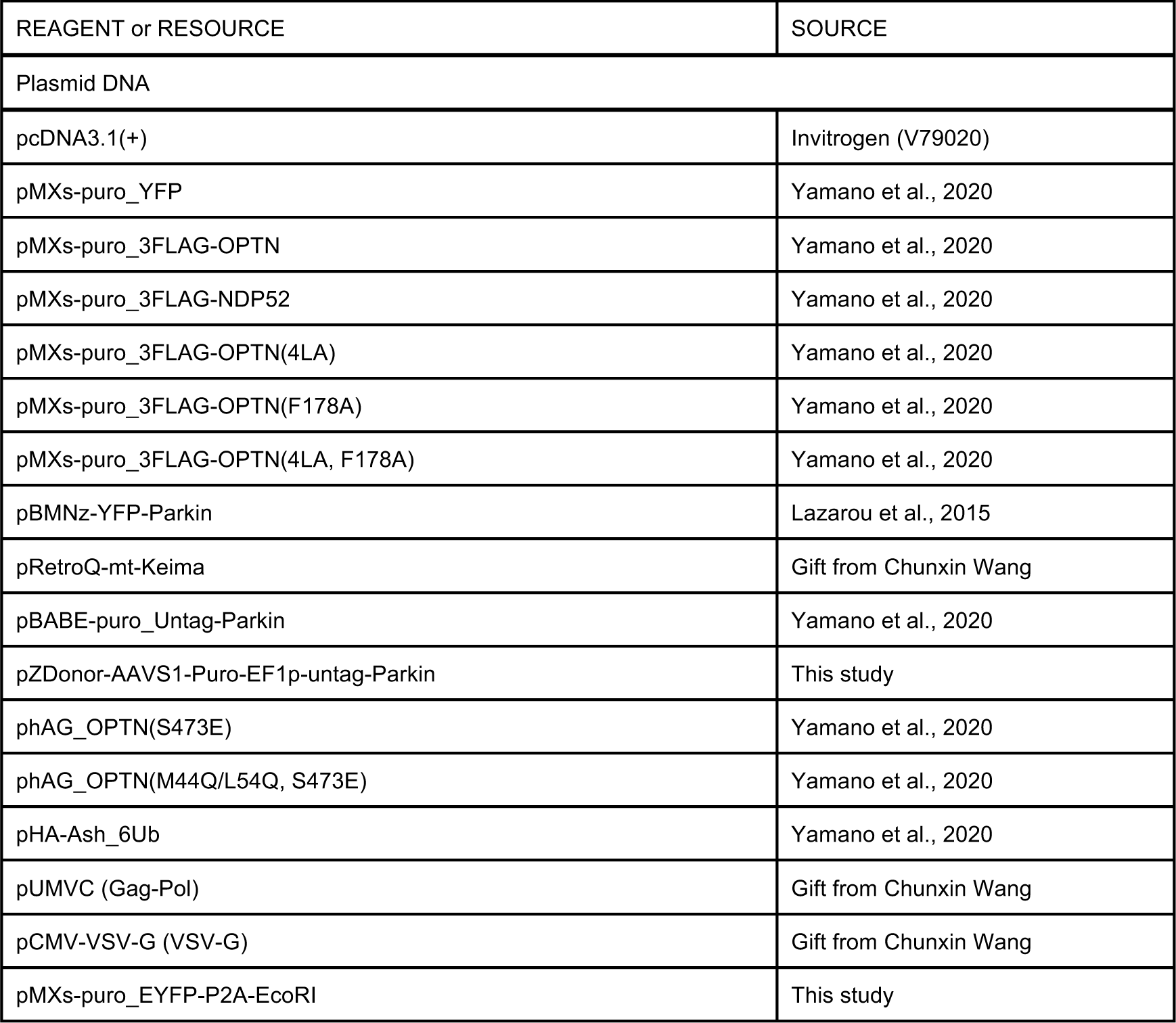
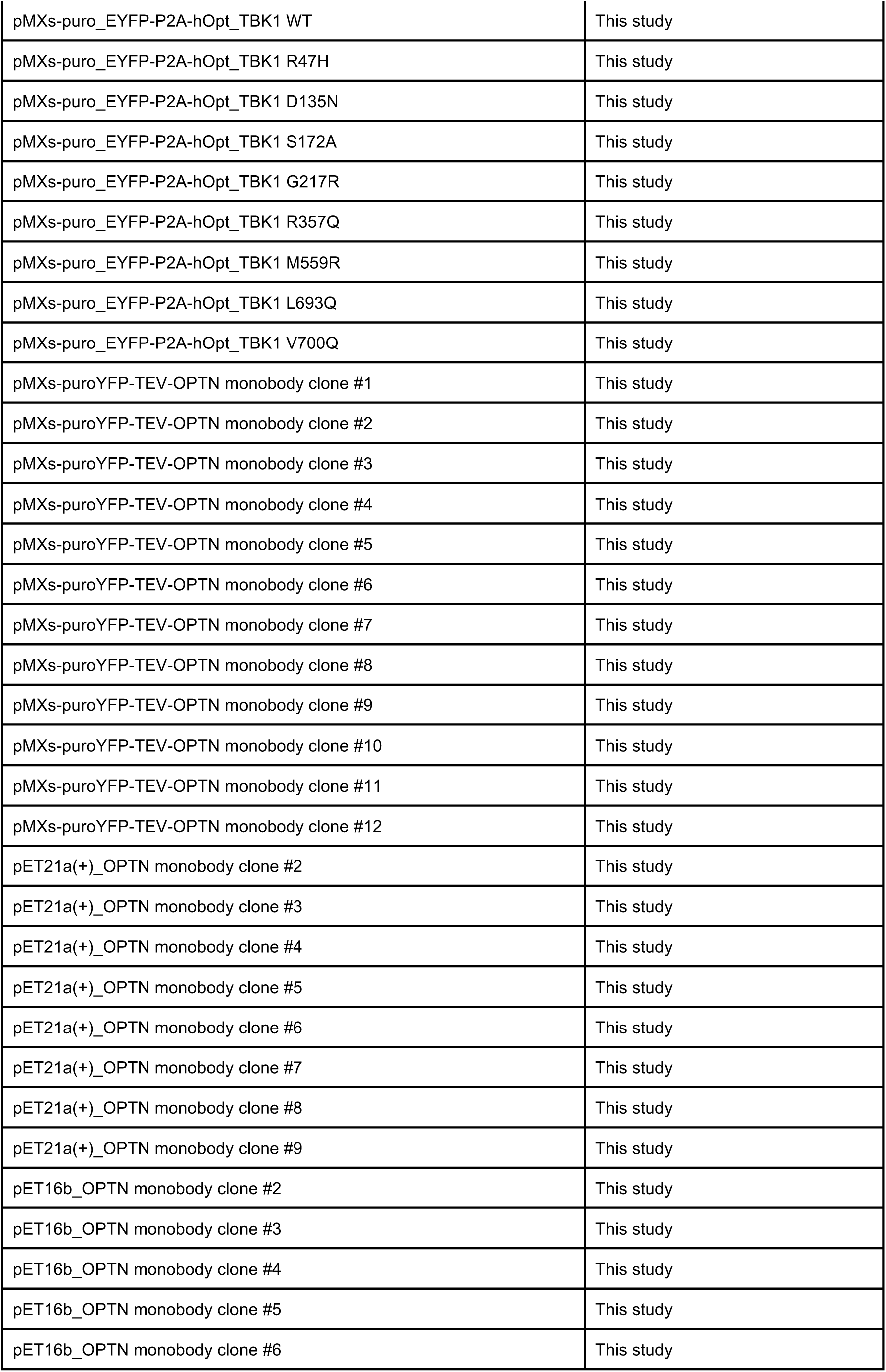
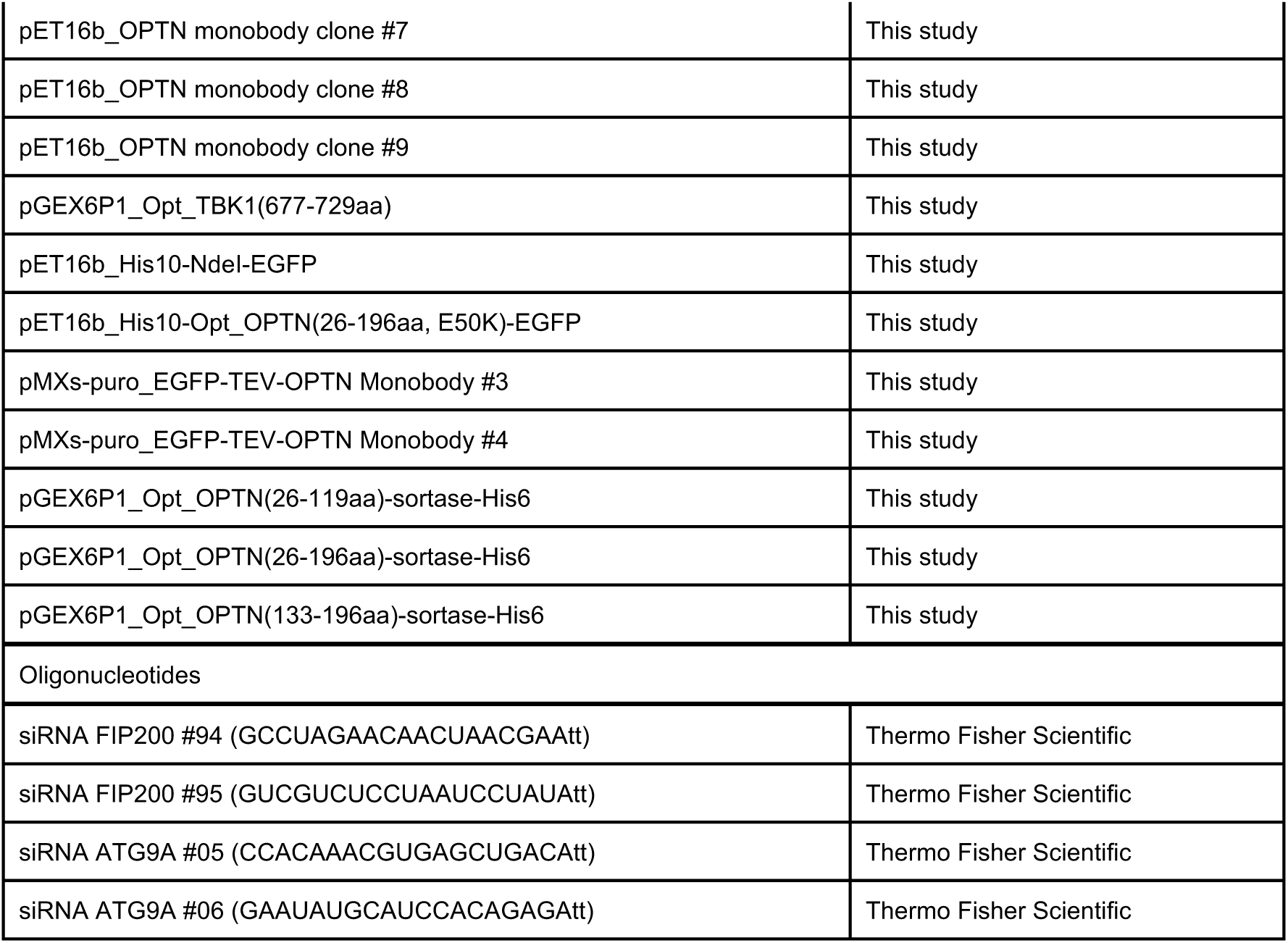
Table for Plasmids and siRNAs

## Acknowledgments

We thank Dr. Richard J. Youle and Dr. Chunxin Wang for the *FIP200* KO, *ATG5* KO, *ATG9A* KO and Penta KO HeLa cells, Dr. Yasushi Saeki’s group for *in vitro* sortase reaction.

This work was supported by JSPS KAKENHI Grants JP18H05500, JP18K06237 and 22H02577 (to K.Y.); by JSPS KAKENHI Grants JP22J00707ZA, JP22K15045ZA (to W.K.); JSPS KAKENHI grant JP19H05712, AMED CREST grant JP23gm1410004h0004, Takeda Science Foundation, Nanken-Kyoten, TMDU, and Joint Usage and Joint Research Programs, Institute of Advanced Medical Sciences, Tokushima University (to N.M.); by JSPS KAKENHI Grants JP22H00419 (to K.T.); by JSPS KAKENHI Grants 19H05287, 21H00278, and JST PRESTO JPMJPR19K6 (to G.H.); by AMED Grants JP21zf0127004 (to H.M.).

## Author contributions

K.Y., M.S., R.K. and W.K. and prepared plasmid materials, constructed stable cell lines, performed the experiments and analyzed the data. K.N., A.S. and G.H. created monobodies and analyzed the data. K.Y. and G.H. designed the research. K.Y. wrote the manuscript with help and supervision from K.T., H.M. and N.M.. The authors declare no competing financial interests.

## Figure Legends

**Supplemental Figure 1 a** Parkin-expressing HeLa cells treated with valinomycin for 1 hr were analyzed by immunostaining with the indicated antibodies. Bars, 10 μm. Schematic representation of autophagy adaptor localization is shown to the right. **b** HeLa cells stably expressing Parkin and GFP-LC3B treated with valinomycin for 1 hr were analyzed by immunostaining with the indicated antibodies. Bars, 10 μm.

**Supplemental Figure 2 a** The indicated proteins in WT, Penta KO, *FIP200* KO, *ATG5* KO, and *ATG9A* KO HeLa cells were analyzed by immunoblotting. **b** Protein levels in (a) were quantified. Protein levels in *FIP200* KO cells were set to 100. Error bars represent mean ± s.d. of three independent experiments.

**Supplemental Figure 3 a** Kinetic parameters of the OPTN monobodies were determined by Bio-layer interferometry. A biotin-labeled OPTN (26-196 aa) was immobilized on a sensor ship. Various concentrations (40, 20, 10, 5, and 2.2 nM) of monobodies (MonoB) #3 and #4 were used for the kinetic analyses. Experimental data are shown in blue and the 1:1 binding model is shown black. **b** Kinetic parameters of the monobodies.

## References

1. Matoba, K. et al. Atg9 is a lipid scramblase that mediates autophagosomal membrane expansion. Nat Struct Mol Biol 27, 1185–1193, doi:10.1038/s41594-020-00518-w (2020).

2. Maeda, S. et al. Structure, lipid scrambling activity and role in autophagosome formation of ATG9A. Nat Struct Mol Biol 27, 1194–1201, doi:10.1038/s41594-020-00520-2 (2020).

3. Sawa-Makarska, J. et al. Reconstitution of autophagosome nucleation defines Atg9 vesicles as seeds for membrane formation. Science 369, doi:10.1126/science.aaz7714 (2020).

4. Yamamoto, H. et al. The Intrinsically Disordered Protein Atg13 Mediates Supramolecular Assembly of Autophagy Initiation Complexes. Dev Cell 38, 86–99, doi:10.1016/j.devcel.2016.06.015 (2016).

5. Fujioka, Y. et al. Phase separation organizes the site of autophagosome formation. Nature 578, 301–305, doi:10.1038/s41586-020-1977-6 (2020).

6. Zhou, C. et al. Regulation of mATG9 trafficking by Src- and ULK1-mediated phosphorylation in basal and starvation-induced autophagy. Cell Res 27, 184–201, doi:10.1038/cr.2016.146 (2017).

7. Russell, R. C. et al. ULK1 induces autophagy by phosphorylating Beclin-1 and activating VPS34 lipid kinase. Nat Cell Biol 15, 741–750, doi:10.1038/ncb2757 (2013).

8. Wold, M. S., Lim, J., Lachance, V., Deng, Z. & Yue, Z. ULK1-mediated phosphorylation of ATG14 promotes autophagy and is impaired in Huntington’s disease models. Mol Neurodegener 11, 76, doi:10.1186/s13024-016-0141-0 (2016).

9. Park, J. M. et al. The ULK1 complex mediates MTORC1 signaling to the autophagy initiation machinery via binding and phosphorylating ATG14. Autophagy 12, 547–564, doi:10.1080/15548627.2016.1140293 (2016).

10. Egan, D. F. et al. Small Molecule Inhibition of the Autophagy Kinase ULK1 and Identification of ULK1 Substrates. Mol Cell 59, 285–297, doi:10.1016/j.molcel.2015.05.031 (2015).

11. Obara, K., Sekito, T., Niimi, K. & Ohsumi, Y. The Atg18-Atg2 complex is recruited to autophagic membranes via phosphatidylinositol 3-phosphate and exerts an essential function. J Biol Chem 283, 23972–23980, doi:10.1074/jbc.M803180200 (2008).

12. Tsuboyama, K. et al. The ATG conjugation systems are important for degradation of the inner autophagosomal membrane. Science 354, 1036–1041, doi:10.1126/science.aaf6136 (2016).

13. Kabeya, Y. et al. LC3, a mammalian homologue of yeast Apg8p, is localized in autophagosome membranes after processing. EMBO J 19, 5720–5728, doi:10.1093/emboj/19.21.5720 (2000).

14. Nguyen, T. N. et al. Atg8 family LC3/GABARAP proteins are crucial for autophagosome-lysosome fusion but not autophagosome formation during PINK1/Parkin mitophagy and starvation. J Cell Biol 215, 857–874, doi:10.1083/jcb.201607039 (2016).

15. Jung, C. H. et al. ULK-Atg13-FIP200 complexes mediate mTOR signaling to the autophagy machinery. Mol Biol Cell 20, 1992–2003, doi:10.1091/mbc.E08-12-1249 (2009).

16. Hosokawa, N. et al. Nutrient-dependent mTORC1 association with the ULK1-Atg13-FIP200 complex required for autophagy. Mol Biol Cell 20, 1981–1991, doi:10.1091/mbc.E08-12-1248 (2009).

17. Vargas, J. N. S., Hamasaki, M., Kawabata, T., Youle, R. J. & Yoshimori, T. The mechanisms and roles of selective autophagy in mammals. Nat Rev Mol Cell Biol, doi:10.1038/s41580-022-00542-2 (2022).

18. Yamano, K., Matsuda, N. & Tanaka, K. The ubiquitin signal and autophagy: an orchestrated dance leading to mitochondrial degradation. EMBO Rep 17, 300–316, doi:10.15252/embr.201541486 (2016).

19. Onishi, M., Yamano, K., Sato, M., Matsuda, N. & Okamoto, K. Molecular mechanisms and physiological functions of mitophagy. EMBO J 40, e104705, doi:10.15252/embj.2020104705 (2021).

20. Pickles, S., Vigie, P. & Youle, R. J. Mitophagy and Quality Control Mechanisms in Mitochondrial Maintenance. Curr Biol 28, R170–R185, doi:10.1016/j.cub.2018.01.004 (2018).

21. Kitada, T. et al. Mutations in the parkin gene cause autosomal recessive juvenile parkinsonism. Nature 392, 605–608, doi:10.1038/33416 (1998).

22. Valente, E. M. et al. Hereditary early-onset Parkinson’s disease caused by mutations in PINK1. Science 304, 1158–1160, doi:10.1126/science.1096284 (2004).

23. Jin, S. M. et al. Mitochondrial membrane potential regulates PINK1 import and proteolytic destabilization by PARL. J Cell Biol 191, 933–942, doi:10.1083/jcb.201008084 (2010).

24. Deas, E. et al. PINK1 cleavage at position A103 by the mitochondrial protease PARL. Hum Mol Genet 20, 867–879, doi:10.1093/hmg/ddq526 (2011).

25. Shi, G. et al. Functional alteration of PARL contributes to mitochondrial dysregulation in Parkinson’s disease. Hum Mol Genet 20, 1966–1974, doi:10.1093/hmg/ddr077 (2011).

26. Meissner, C., Lorenz, H., Weihofen, A., Selkoe, D. J. & Lemberg, M. K. The mitochondrial intramembrane protease PARL cleaves human Pink1 to regulate Pink1 trafficking. J Neurochem 117, 856–867, doi:10.1111/j.1471-4159.2011.07253.x (2011).

27. Greene, A. W. et al. Mitochondrial processing peptidase regulates PINK1 processing, import and Parkin recruitment. EMBO Rep 13, 378–385, doi:10.1038/embor.2012.14 (2012).

28. Yamano, K. & Youle, R. J. PINK1 is degraded through the N-end rule pathway. Autophagy 9, 1758–1769, doi:10.4161/auto.24633 (2013).

29. Sekine, S. et al. Reciprocal Roles of Tom7 and OMA1 during Mitochondrial Import and Activation of PINK1. Mol Cell 73, 1028–1043 e1025, doi:10.1016/j.molcel.2019.01.002 (2019).

30. Lazarou, M., Jin, S. M., Kane, L. A. & Youle, R. J. Role of PINK1 binding to the TOM complex and alternate intracellular membranes in recruitment and activation of the E3 ligase Parkin. Dev Cell 22, 320–333, doi:10.1016/j.devcel.2011.12.014 (2012).

31. Hasson, S. A. et al. High-content genome-wide RNAi screens identify regulators of parkin upstream of mitophagy. Nature 504, 291–295, doi:10.1038/nature12748 (2013).

32. Okatsu, K. et al. A dimeric PINK1-containing complex on depolarized mitochondria stimulates Parkin recruitment. J Biol Chem 288, 36372–36384, doi:10.1074/jbc.M113.509653 (2013).

33. Matsuda, N. et al. PINK1 stabilized by mitochondrial depolarization recruits Parkin to damaged mitochondria and activates latent Parkin for mitophagy. J Cell Biol 189, 211–221, doi:10.1083/jcb.200910140 (2010).

34. Narendra, D., Tanaka, A., Suen, D. F. & Youle, R. J. Parkin is recruited selectively to impaired mitochondria and promotes their autophagy. J Cell Biol 183, 795–803, doi:10.1083/jcb.200809125 (2008).

35. Narendra, D. P. et al. PINK1 is selectively stabilized on impaired mitochondria to activate Parkin. PLoS Biol 8, e1000298, doi:10.1371/journal.pbio.1000298 (2010).

36. Shiba-Fukushima, K. et al. PINK1-mediated phosphorylation of the Parkin ubiquitin-like domain primes mitochondrial translocation of Parkin and regulates mitophagy. Sci Rep 2, 1002, doi:10.1038/srep01002 (2012).

37. Kondapalli, C. et al. PINK1 is activated by mitochondrial membrane potential depolarization and stimulates Parkin E3 ligase activity by phosphorylating Serine 65. Open Biol 2, 120080, doi:10.1098/rsob.120080 (2012).

38. Koyano, F. et al. Ubiquitin is phosphorylated by PINK1 to activate parkin. Nature 510, 162–166, doi:10.1038/nature13392 (2014).

39. Kazlauskaite, A. et al. Parkin is activated by PINK1-dependent phosphorylation of ubiquitin at Ser65. Biochem J 460, 127–139, doi:10.1042/BJ20140334 (2014).

40. Kane, L. A. et al. PINK1 phosphorylates ubiquitin to activate Parkin E3 ubiquitin ligase activity. J Cell Biol 205, 143–153, doi:10.1083/jcb.201402104 (2014).

41. Gundogdu, M., Tadayon, R., Salzano, G., Shaw, G. S. & Walden, H. A mechanistic review of Parkin activation. Biochim Biophys Acta Gen Subj 1865, 129894, doi:10.1016/j.bbagen.2021.129894 (2021).

42. Yamano, K. et al. Site-specific Interaction Mapping of Phosphorylated Ubiquitin to Uncover Parkin Activation. J Biol Chem 290, 25199–25211, doi:10.1074/jbc.M115.671446 (2015).

43. Gladkova, C., Maslen, S. L., Skehel, J. M. & Komander, D. Mechanism of parkin activation by PINK1. Nature 559, 410–414, doi:10.1038/s41586-018-0224-x (2018).

44. Sauve, V. et al. Mechanism of parkin activation by phosphorylation. Nat Struct Mol Biol 25, 623–630, doi:10.1038/s41594-018-0088-7 (2018).

45. Ordureau, A. et al. Quantitative proteomics reveal a feedforward mechanism for mitochondrial PARKIN translocation and ubiquitin chain synthesis. Mol Cell 56, 360–375, doi:10.1016/j.molcel.2014.09.007 (2014).

46. Okatsu, K. et al. Phosphorylated ubiquitin chain is the genuine Parkin receptor. J Cell Biol 209, 111–128, doi:10.1083/jcb.201410050 (2015).

47. Ordureau, A. et al. Defining roles of PARKIN and ubiquitin phosphorylation by PINK1 in mitochondrial quality control using a ubiquitin replacement strategy. Proc Natl Acad Sci U S A 112, 6637–6642, doi:10.1073/pnas.1506593112 (2015).

48. Hayashida, R. et al. Elucidation of ubiquitin-conjugating enzymes that interact with RBR-type ubiquitin ligases using a liquid-liquid phase separation-based method. J Biol Chem 299, 102822, doi:10.1016/j.jbc.2022.102822 (2022).

49. Lazarou, M. et al. The ubiquitin kinase PINK1 recruits autophagy receptors to induce mitophagy. Nature 524, 309–314, doi:10.1038/nature14893 (2015).

50. Yamano, K. & Kojima, W. Molecular functions of autophagy adaptors upon ubiquitin-driven mitophagy. Biochim Biophys Acta Gen Subj 1865, 129972, doi:10.1016/j.bbagen.2021.129972 (2021).

51. Adriaenssens, E., Ferrari, L. & Martens, S. Orchestration of selective autophagy by cargo receptors. Curr Biol 32, R1357–R1371, doi:10.1016/j.cub.2022.11.002 (2022).

52. Fu, T. et al. Structural and biochemical advances on the recruitment of the autophagy-initiating ULK and TBK1 complexes by autophagy receptor NDP52. Sci Adv 7, doi:10.1126/sciadv.abi6582 (2021).

53. Shi, X., Chang, C., Yokom, A. L., Jensen, L. E. & Hurley, J. H. The autophagy adaptor NDP52 and the FIP200 coiled-coil allosterically activate ULK1 complex membrane recruitment. Elife 9, doi:10.7554/eLife.59099 (2020).

54. Ravenhill, B. J. et al. The Cargo Receptor NDP52 Initiates Selective Autophagy by Recruiting the ULK Complex to Cytosol-Invading Bacteria. Mol Cell 74, 320–329 e326, doi:10.1016/j.molcel.2019.01.041 (2019).

55. Vargas, J. N. S. et al. Spatiotemporal Control of ULK1 Activation by NDP52 and TBK1 during Selective Autophagy. Mol Cell 74, 347–362 e346, doi:10.1016/j.molcel.2019.02.010 (2019).

56. Yamano, K. et al. Critical role of mitochondrial ubiquitination and the OPTN-ATG9A axis in mitophagy. J Cell Biol 219, doi:10.1083/jcb.201912144 (2020).

57. Zhou, Z. et al. Phosphorylation regulates the binding of autophagy receptors to FIP200 Claw domain for selective autophagy initiation. Nat Commun 12, 1570, doi:10.1038/s41467-021-21874-1 (2021).

58. Wild, P. et al. Phosphorylation of the autophagy receptor optineurin restricts Salmonella growth. Science 333, 228–233, doi:10.1126/science.1205405 (2011).

59. Li, F. et al. Structural insights into the interaction and disease mechanism of neurodegenerative disease-associated optineurin and TBK1 proteins. Nat Commun 7, 12708, doi:10.1038/ncomms12708 (2016).

60. Thurston, T. L., Ryzhakov, G., Bloor, S., von Muhlinen, N. & Randow, F. The TBK1 adaptor and autophagy receptor NDP52 restricts the proliferation of ubiquitin-coated bacteria. Nat Immunol 10, 1215–1221, doi:10.1038/ni.1800 (2009).

61. Zhang, C. et al. Structural basis of STING binding with and phosphorylation by TBK1. Nature 567, 394–398, doi:10.1038/s41586-019-1000-2 (2019).

62. Heo, J. M., Ordureau, A., Paulo, J. A., Rinehart, J. & Harper, J. W. The PINK1-PARKIN Mitochondrial Ubiquitylation Pathway Drives a Program of OPTN/NDP52 Recruitment and TBK1 Activation to Promote Mitophagy. Mol Cell 60, 7–20, doi:10.1016/j.molcel.2015.08.016 (2015).

63. Richter, B. et al. Phosphorylation of OPTN by TBK1 enhances its binding to Ub chains and promotes selective autophagy of damaged mitochondria. Proc Natl Acad Sci U S A 113, 4039–4044, doi:10.1073/pnas.1523926113 (2016).

64. Heo, J. M. et al. RAB7A phosphorylation by TBK1 promotes mitophagy via the PINK-PARKIN pathway. Sci Adv 4, eaav0443, doi:10.1126/sciadv.aav0443 (2018).

65. Herhaus, L. et al. TBK1-mediated phosphorylation of LC3C and GABARAP-L2 controls autophagosome shedding by ATG4 protease. EMBO Rep 21, e48317, doi:10.15252/embr.201948317 (2020).

66. Cirulli, E. T. et al. Exome sequencing in amyotrophic lateral sclerosis identifies risk genes and pathways. Science 347, 1436–1441, doi:10.1126/science.aaa3650 (2015).

67. Evans, C. S. & Holzbaur, E. L. F. Autophagy and mitophagy in ALS. Neurobiol Dis 122, 35–40, doi:10.1016/j.nbd.2018.07.005 (2019).

68. Kachaner, D. et al. Plk1-dependent phosphorylation of optineurin provides a negative feedback mechanism for mitotic progression. Mol Cell 45, 553–566, doi:10.1016/j.molcel.2011.12.030 (2012).

69. Padman, B. S. et al. LC3/GABARAPs drive ubiquitin-independent recruitment of Optineurin and NDP52 to amplify mitophagy. Nat Commun 10, 408, doi:10.1038/s41467-019-08335-6 (2019).

70. Johansen, T. & Lamark, T. Selective Autophagy: ATG8 Family Proteins, LIR Motifs and Cargo Receptors. J Mol Biol 432, 80–103, doi:10.1016/j.jmb.2019.07.016 (2020).

71. Swatek, K. N. & Komander, D. Ubiquitin modifications. Cell Res 26, 399–422, doi:10.1038/cr.2016.39 (2016).

72. Watanabe, T. et al. Genetic visualization of protein interactions harnessing liquid phase transitions. Sci Rep 7, 46380, doi:10.1038/srep46380 (2017).

73. Ye, J. et al. Effects of ALS-associated TANK binding kinase 1 mutations on protein-protein interactions and kinase activity. Proc Natl Acad Sci U S A 116, 24517–24526, doi:10.1073/pnas.1915732116 (2019).

74. Harding, O. et al. ALS- and FTD-associated missense mutations in TBK1 differentially disrupt mitophagy. Proc Natl Acad Sci U S A 118, doi:10.1073/pnas.2025053118 (2021).

75. Katayama, H., Kogure, T., Mizushima, N., Yoshimori, T. & Miyawaki, A. A sensitive and quantitative technique for detecting autophagic events based on lysosomal delivery. Chem Biol 18, 1042–1052, doi:10.1016/j.chembiol.2011.05.013 (2011).

76. Koide, A., Bailey, C. W., Huang, X. & Koide, S. The fibronectin type III domain as a scaffold for novel binding proteins. J Mol Biol 284, 1141–1151, doi:10.1006/jmbi.1998.2238 (1998).

77. Kondo, T. et al. Antibody-like proteins that capture and neutralize SARS-CoV-2. Sci Adv 6, doi:10.1126/sciadv.abd3916 (2020).

78. Alam, M., Hasan, G. M. & Hassan, M. I. A review on the role of TANK-binding kinase 1 signaling in cancer. Int J Biol Macromol 183, 2364–2375, doi:10.1016/j.ijbiomac.2021.06.022 (2021).

79. Helgason, E., Phung, Q. T. & Dueber, E. C. Recent insights into the complexity of Tank-binding kinase 1 signaling networks: the emerging role of cellular localization in the activation and substrate specificity of TBK1. FEBS Lett 587, 1230–1237, doi:10.1016/j.febslet.2013.01.059 (2013).

80. Shang, G., Zhang, C., Chen, Z. J., Bai, X. C. & Zhang, X. Cryo-EM structures of STING reveal its mechanism of activation by cyclic GMP-AMP. Nature 567, 389–393, doi:10.1038/s41586-019-0998-5 (2019).

81. Cai, X., Chiu, Y. H. & Chen, Z. J. The cGAS-cGAMP-STING pathway of cytosolic DNA sensing and signaling. Mol Cell 54, 289–296, doi:10.1016/j.molcel.2014.03.040 (2014).

82. Rezaie, T. et al. Adult-onset primary open-angle glaucoma caused by mutations in optineurin. Science 295, 1077–1079, doi:10.1126/science.1066901 (2002).

83. Minegishi, Y. et al. Enhanced optineurin E50K-TBK1 interaction evokes protein insolubility and initiates familial primary open-angle glaucoma. Hum Mol Genet 22, 3559–3567, doi:10.1093/hmg/ddt210 (2013).

84. Nezich, C. L., Wang, C., Fogel, A. I. & Youle, R. J. MiT/TFE transcription factors are activated during mitophagy downstream of Parkin and Atg5. J Cell Biol 210, 435–450, doi:10.1083/jcb.201501002 (2015).

