## Supplementary figures and images for "Optineurin provides a mitophagy contact site for TBK1 activation"

### Supplementary Figure 1

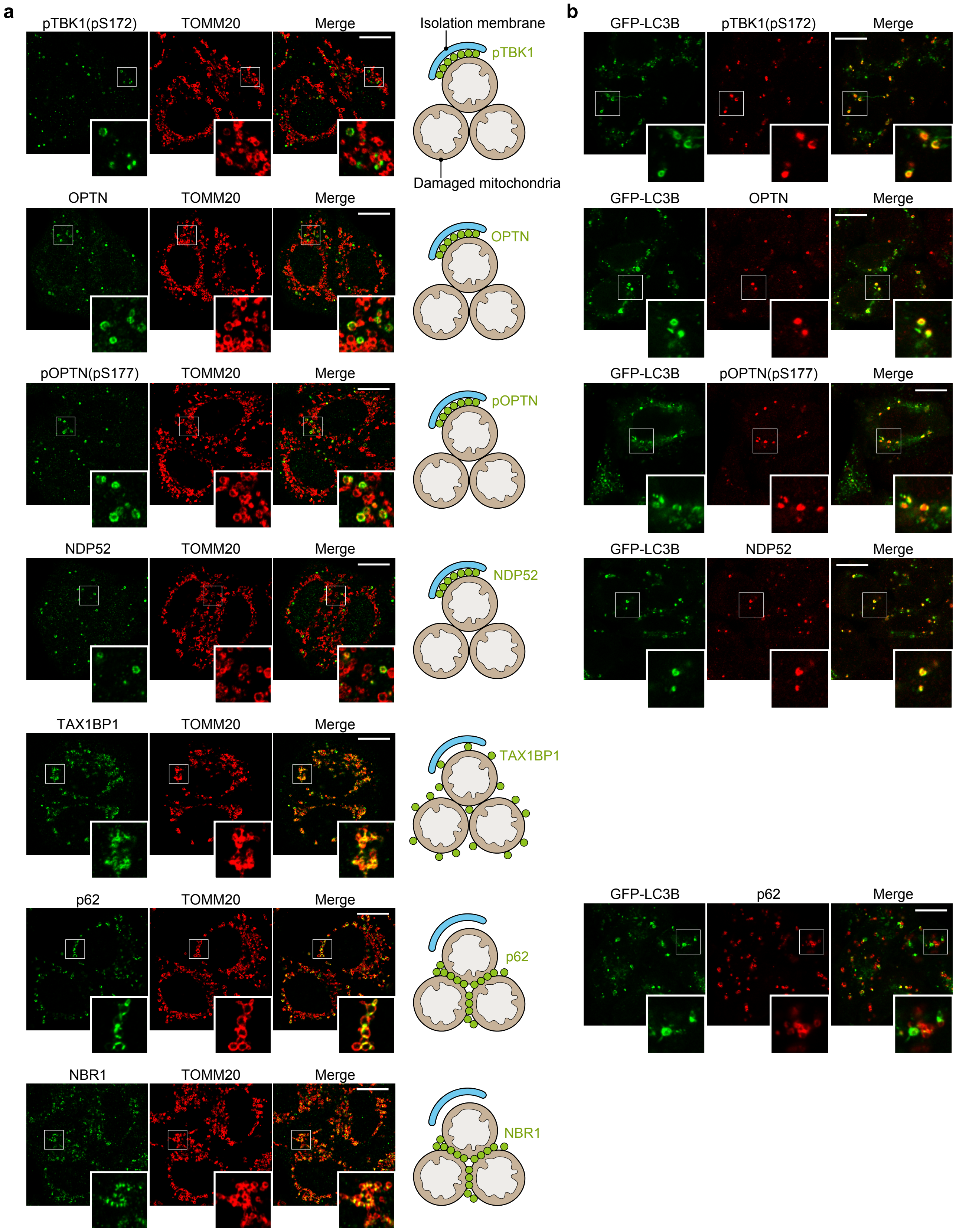


Yamano et al Supplementary Figure 1

### Supplementary Figure 2

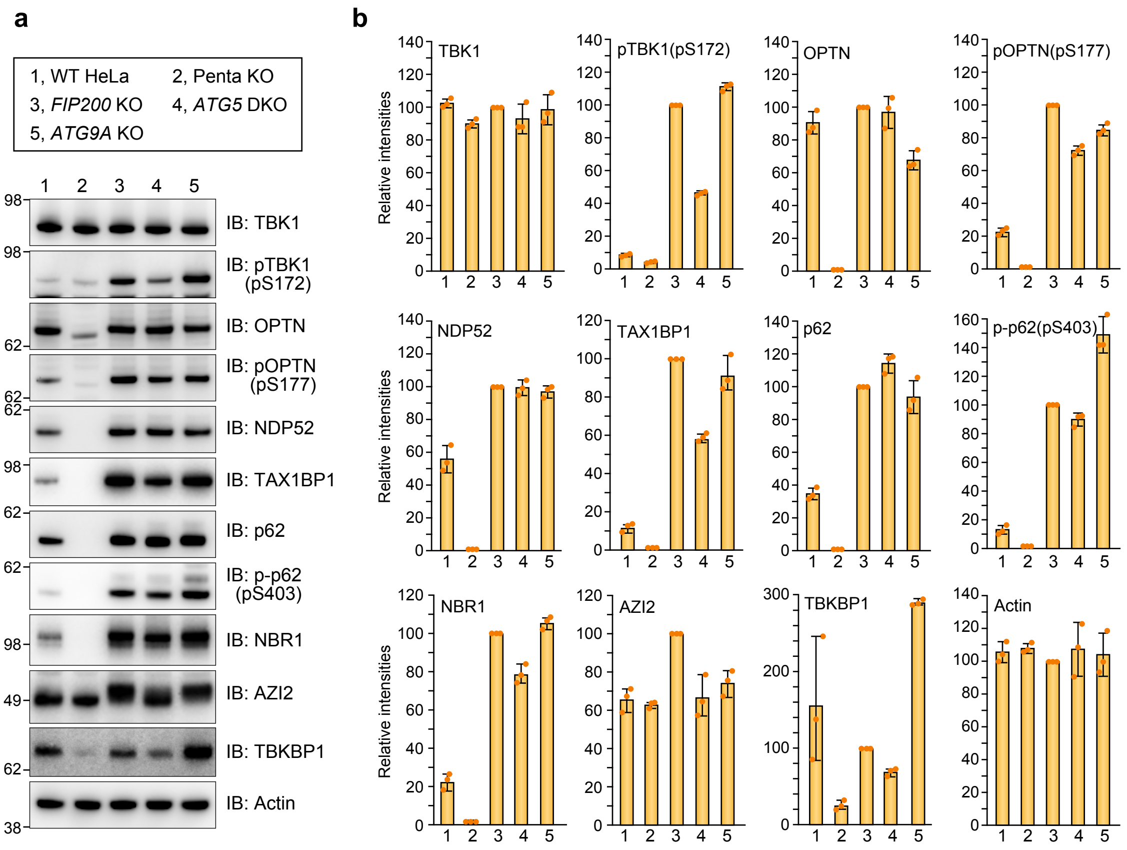


Yamano et al Supplementary Figure 2

### Supplementary Figure 3

**a**

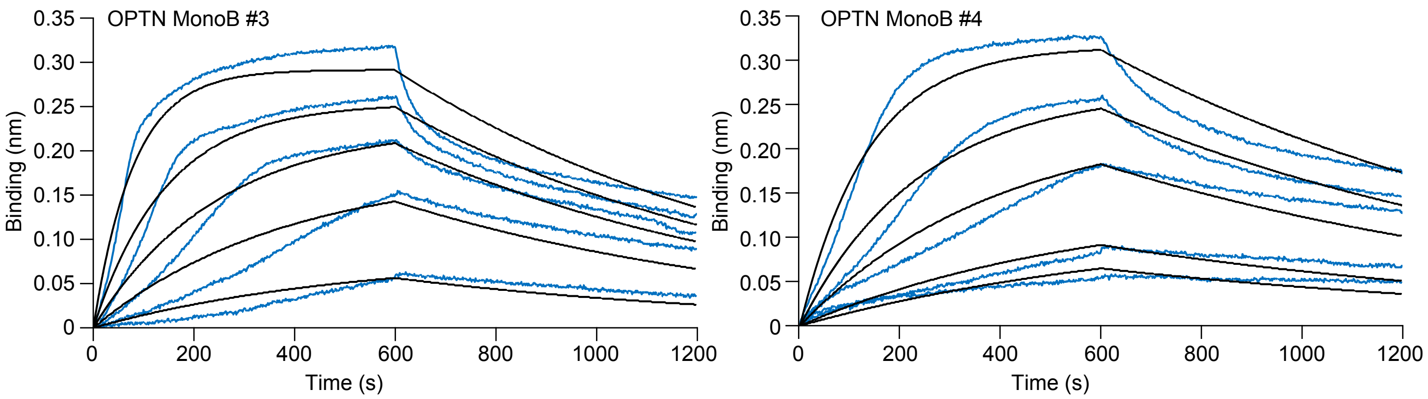


**b**

| Names | $K_D$ (nM) | $k_{on}$ (1/Ms) ×$10^4$ | $k_{off}$ (1/s) ×$10^{-4}$ |
|---|---|---|---|
| OPTN MonoB #3 | 4.56 | 27.81 | 12.69 |
| OPTN MonoB #4 | 6.19 | 15.84 | 9.81 |

Yamano et al Supplementary Figure 3
